# Muscle-controlled physics simulations of the emu (a large running bird) resolve grounded running paradox

**DOI:** 10.1101/2024.01.17.575928

**Authors:** Pasha A. van Bijlert, A.J. “Knoek” van Soest, Anne S. Schulp, Karl T. Bates

## Abstract

Humans and birds utilize very different running styles. Unlike humans, birds adopt “grounded running” at intermediate speeds – a running gait where at least one foot is always in contact with the ground. Avian grounded running is paradoxical: animals tend to minimize locomotor energy expenditure, but birds prefer grounded running despite incurring higher energy costs. Using predictive gait simulations of the emu (*Dromaius novaehollandiae*), we resolve this paradox by demonstrating that grounded running represents an energetic optimum for birds. Our virtual experiments decoupled biomechanically relevant anatomical features that cannot be isolated in a real bird. The avian body plan prevents (near) vertical leg postures while running, making the running style used by humans impossible. Under this anatomical constraint, grounded running is optimal if the muscles produce the highest forces in crouched postures, as is true in most birds. Anatomical similarities between birds and non-avian dinosaurs suggest that, as a behavior, avian grounded running first evolved within non-avian theropods.

## Introduction

Understanding why animals move in certain ways is a fundamental goal of biomechanics, ecology, and evolutionary biology. It is reasonable to assume that animals evolve functional compromises between different features beneficial to their survival (*1*). In terrestrial vertebrates, many factors have been suggested to affect gait selection, including (minimization of) energy expenditure (*2–6*), center of mass (*COM*) movements (*7*, *8*), neuromuscular factors (*9–11*), gross morphology (*12*, *13*), injury prevention (*14*), and skeletal stresses (*15–17*). By nature, some of these represent conflicting demands, and understanding which factors are dominant in an organism can shed light on what selective pressures may have shaped its evolutionary history.

With such fundamental tradeoffs in mind, bipedal walking and running (striding) gaits present some clear mechanical challenges: life on two legs is less stable than on four, and stresses on the limbs are roughly twice as high. Despite these apparent challenges, large ratite birds are notable for their ability to reach speeds up to 14 – 17 m s^-1^ (50 – 60 km h^-1^) (*18–22*), and both ratites and humans are capable of economical locomotion (*3*, *6*). It has been well-established that animals prefer different gaits depending on the desired speed, which contributes to minimization of the metabolic cost of transport (*MCOT*, in J kg^-1^ m^-1^) (*2*, *5*, *6*).

In bipeds, the forward speed of the *COM* is minimal near midstance, regardless of the gait. The vertical position of the *COM* at this instant is gait dependent: when walking, the *COM* is at its highest near midstance, whereas when running the *COM* is at its lowest near midstance (Figure 1 A). This observation can be described in energetic terms: walking is a pendular (vaulting) gait, where kinetic (*E*_K_) and potential energies (*E*_P_) of the *COM* are out-of-phase (*1*, *7*, *8*, *13*, *23*, *24*). In contrast, fluctuations of *COM* energy are in-phase during running, resembling a bouncing gait (*1*, *7*, *8*, *13*, *24*).

**Figure 1.**
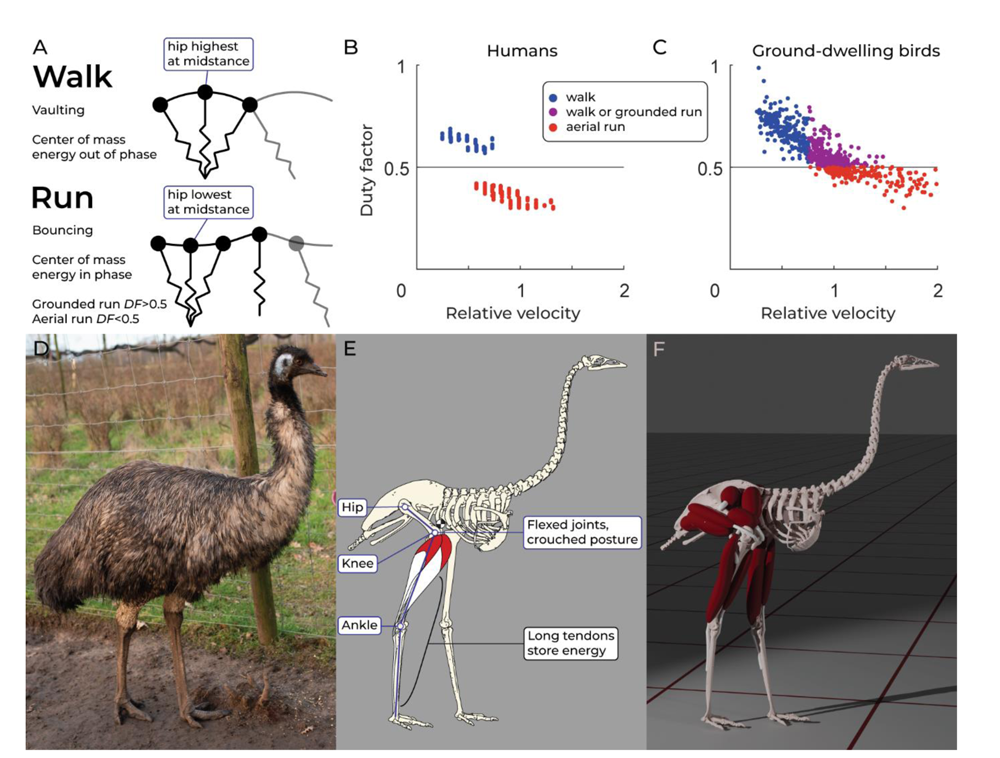
Features of walking and running in birds and humans focused on in this study. (**A**) When walking, the hip reaches its highest point during midstance (kinetic and potential energy of the center of mass are out of phase). When running, the hip reaches its lowest point at midstance (center of mass energies are in-phase). Running gaits are further categorized by duty factor (*DF*): in grounded running, duty factor is above 0.5 (and there is thus no aerial phase). In aerial running, duty factor is below 0.5 (signifying an aerial phase during the gait cycle). (**B**) and (**C**) Duty factor plotted against relative velocity (dimensionless speed, see methods) in humans and birds, respectively. In humans, the walk to run transition is marked by a stark reduction in duty factor. In contrast, birds first smoothly transition to grounded running, and thus their gaits cannot be recognized based on duty factor alone. It is currently unclear why birds prefer grounded running at intermediate speeds. Data plotted from Bishop et al. 2018. (**D**) In the emu (*Dromaius novaehollandiae*), as in most birds, the hip and knee joints are enveloped in feathers, obscuring the fact that (**E**) most birds habitually keep their three functional leg segments in crouched postures, because their muscles are strongest near these postures. Grounded running in birds may be related to these crouched postures, and/or the presence of extraordinarily long tendons in the distal hindlimb, which enable elastic energy storage. A fully extended posture is impossible for birds due to the forward placement of the center of mass (checkered circle). (**F**) Our musculoskeletal model of the emu, developed for this study, enabled us to decouple the effects of posture and tendon elastic storage on running gaits.

In humans, the walk-to-run transition is accompanied by an abrupt drop in duty factor (*DF*, fraction of the stride period a foot is in contact with the ground) (Figure 1 B) (*25*, *26*). Above the transition speed, humans switch to a running gait with an aerial phase (*DF* < 0.5). In birds, no such abrupt transition in stride kinematics exists (*5*, *13*, *25–28*). Birds transition from walking to running without a discontinuity in *DF* (Figure 1 C): they first switch to what has become known as a grounded running gait; that is a gait with no aerial phase (*DF* > 0.5), but with in-phase *COM* oscillations (*5*, *13*, *25*, *27*, *28*). At higher speeds, *DF* steadily decreases until aerial running occurs. This means that the absence of an aerial phase is not enough to determine whether birds (and indeed, even many quadrupedal animals) are running or not (*7*) (Figure 1 C).

Grounded running has higher energy costs than aerial running, both at the same speed (*24*) and when the gaits are compared at different speeds (*5*, *6*, *28*). In humans, this is often attributed to the higher net joint moments required in crouched postures (*24*, *29*). Avian preference for grounded running at intermediate speeds is thus paradoxical: they appear to habitually prefer a running style that is energetically costly, whereas animals usually adopt energetically (near-) optimal gaits. This paradox has been the subject of much debate (*5*, *6*, *20*, *25*, *28*, *30*, *31*). Grounded running decreases accelerations at the head, increases stability, and may aid injury prevention (*24*, *25*, *29*, *31*). The “walk to run” transition in ostriches occurs at a higher speed than the “run to walk” transition (*20*). While these aforementioned findings suggest the possibility that energetics are not the most important determinant in avian gait selection, this seemingly contradicts that birds do minimize *MCOT* during gait selection (*5*, *6*). Alternatively, there may be anatomical reasons why grounded running is metabolically optimal for birds, but is suboptimal in humans.

One possible anatomical reason for the avian preference for grounded running is elastic energy storage in tendons (*5*, *20*). Locomotor adaptations in large ratite birds are well established: they have very long digital flexor tendons that store energy when the toes are extended (Figure 1 E) (*18*, *19*, *32–34*), and similar mechanisms have been demonstrated in much smaller birds as well (*9*, *35*). It has been estimated that elastic energy storage is more than twice as high in ostriches than in humans (*34*), and representing an important factor in avian gait selection.

Another important anatomical feature related to grounded running is the habitually crouched (i.e., flexed) postures that birds adopt (*13*, *25–27*, *36*) (Figure 1 D,E). Animals tend to adopt joint poses at which their muscles can generate the highest forces (*10*), and tendon elasticity in birds biases them towards more crouched postures (*37*). Humans are an exception in this regard, preferring a fully extended, columnar posture (*36*), even though they are strongest in crouched postures (*38*). Such a fully extended posture is impossible for birds, because their *COM* lies in front of the hip joint, requiring crouched postures to maintain balance (*36*, *39*, *40*) (Figure 1 E and 2 B). Interestingly, crouched postures appear to be a prerequisite for grounded running in humans (*24*, *29*). This is somewhat supported by recent work modelling bird locomotion as a simple spring-loaded inverted pendulum (SLIP) (Figure 1A), suggesting that grounded running requires sufficient spring compliance (*30*). The SLIP-model reduces the hindlimb to a rigid rod mounted on a compliant spring. Unfortunately, there is no clear correspondence between changes in SLIP-model leg length and joint poses in animals modelled in (*30*), and it is unclear if energy storage in the SLIP-model spring has a biological interpretation. Thus, the relative contributions of posture and elastic-storage, and therefore the mechanistic triggers for habitual grounded running in birds, remain poorly understood.

Decoupling the mechanical effects of tendon elasticity and crouched postural tunings in a bird in-vivo is impractical (if not impossible) and would raise ethical concerns. Furthermore, birds are not easily trained to adopt different running styles at the same speed, further preventing direct comparisons of different gaits (*5*, *28*). Instead, musculoskeletal modelling represents an ideal vehicle to test such a phenomenon. Multibody dynamic analysis using such models has been used in both human (*41–47*) and comparative (*37*, *39*, *48–53*) contexts, and can provide insights complementary to physical experiments (*13*). When such models are combined with optimal control methods (e.g., (*41*, *54*)), gaits can be found without using kinematic (motion capture) data as an input, often referred to as “predictive simulation”. Predictive simulation is particularly attractive for the present study: it enables us to investigate how (altered) musculoskeletal design affects gait selection, without *a priori* biasing the model towards desired (measured) gaits. Such insights would not be possible through experimentation, but requires that ample data exist to validate simulator outputs. The emu (*Dromaius novaehollandiae*) is a large, flightless bird native to Australia (*22*, *55*) (Figure 1 D). Emu locomotion and anatomy are well-studied (*6*, *17*, *19*, *21*, *25–27*, *33*, *48*, *49*, *56–58*), making emus a suitable model organism for locomotor research. In this study, we constructed a new 3D musculoskeletal model of the emu for predictive multibody dynamic gait simulations (Figure 1 F, Figure 2).

**Figure 2.**
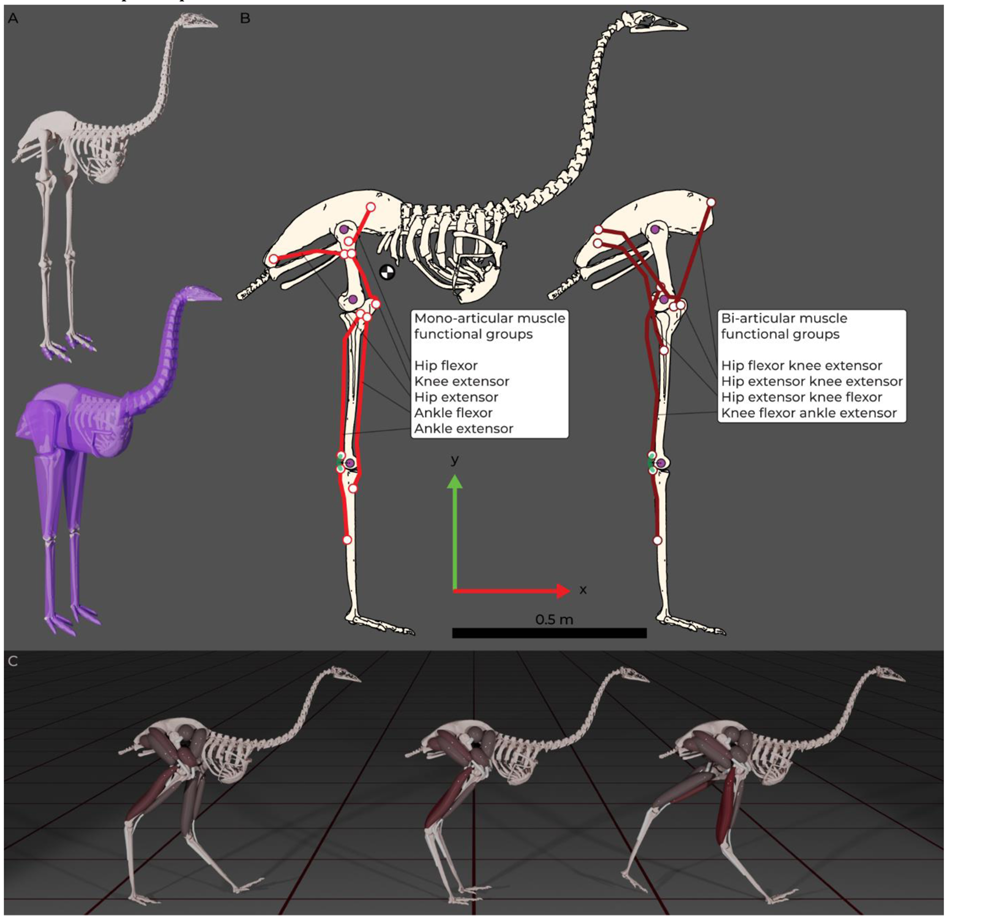
Overview of methodological steps from model construction to dynamic simulation. (**A**). Top: Symmetrized skeletal model derived from CT-scans, positioned in a neutral reference pose. The model has 10 contact spheres per foot. Below: Skin outlines for the model, based on convex-hull reconstructions (**B**). Orthographic projection of the joint centers and paths (lines of action) of the muscle functional groups. Ankle extensors have wrapping cylinders of 0.03 m radius (in green). We assume a right-handed reference frame. (**C**). Muscle-controlled dynamic simulation, generated de-novo without using measured emu kinematics.

Our primary goal was to investigate the grounded running paradox: *why do birds adopt grounded running gaits at intermediate speeds* (Figure 1 C), *even though studies show* (*24*, *28*) *that this is energetically costly?* We hypothesized that grounded running is an attractive running style for birds because their muscles function optimally in crouched postures, and fully extended (human-like) postures are impossible due to their anatomy. Because grounded running is often referred to as a “compliant gait” with spring-like leg behavior, a secondary goal was to investigate whether grounded running requires any compliance (i.e., any elastic energy storage). Our simulations provided strong support for our over-arching hypothesis and revealed a fundamental tradeoff in gait selection. We found that grounded running is purely an effect of changes in effective leg length – no energy storage (or leg spring compliance) is required, it is an effect of limb posture. Grounded running requires crouched postures to achieve (*24*, *25*, *30*), which results in higher muscle forces (and thus higher energy costs) than columnar postures. However, grounded running results in lower peak ground reaction forces, which decreases energy costs (*4*, *59*). There is thus a tradeoff between minimizing peak muscle forces (columnar postures), and minimizing peak ground reaction forces (crouched postures, grounded running), and humans adopt the former. In contrast, our results demonstrate for the first time that grounded running represents a metabolically optimal gait for crouched bipeds (such as birds). A columnar, human-like running style could have lower energy costs for birds, but this is impossible due to the forward-placement of their center of mass, combined with muscles that generate the most force in crouched postures. As a result, grounded running is energetically optimal at intermediate speeds. Thus, we argue that the paradox in avian gait selection does not exist, and that secondary benefits of grounded running in birds are not at odds with the well-established principle of *MCOT* minimization.

## Results

Our analysis relied on systematically changing the muscular anatomy of the model, and simulating how this affected running styles found through optimization. We hypothesized that grounded running is optimal for bipeds with crouched habitual postures (such as birds), and not necessarily dependent on elastic storage in tendons. To demonstrate this, our simulations decoupled the effects of leg postural tuning and tendon elasticity. Grounded running occurred more often in our model variants if they had crouched tunings, using a variety of different optimization approaches and model formulations. Elastic tendons, an important factor in bird anatomy, tends to increase the occurrence by affecting leg postures, but elastic storage itself was not required for grounded running. We will first demonstrate that systematic differences in our model variants resulted in systemically different postures and gaits, and general patterns related to these differences. We will compare these model variant gaits to real emu data, to demonstrate that all the variants captured the salient features of emu locomotion. After this, we will present results that directly support our hypothesis.

Unless otherwise specified, we present data from optimizations where we minimized both muscle fatigue and energetic cost (“fatigue and *MCOT*”, see methods). Fatigue was parametrized neural input (excitations cubed, see methods), and energy cost as the metabolic cost of transport (*MCOT*). The fatigue-only and *MCOT-*only optimizations are presented as supplementary data and figures. We have analyzed more than 650 trials (excluding local optima and pilot simulations).

### General patterns

Our muscle tuning procedure successfully biased the model variants as desired to relatively columnar, intermediate, and crouched postures (compare hip heights *h* in Figure 3, left to right). Supplementary Video S2 demonstrates the different walking gaits of the model variants. Crouched variants showed a larger range in effective leg length (*L*_eff_) during the stance phase than more columnar variants, both in relative and absolute sense (Figure 3). All elastic tendon model variants adopted more crouched midstance postures and used a wider joint range during locomotion (resulting in lower *h* and a larger range in *L*_eff_) than the rigid tendon variants (Figure 3, top row versus bottom row).

**Figure 3.**
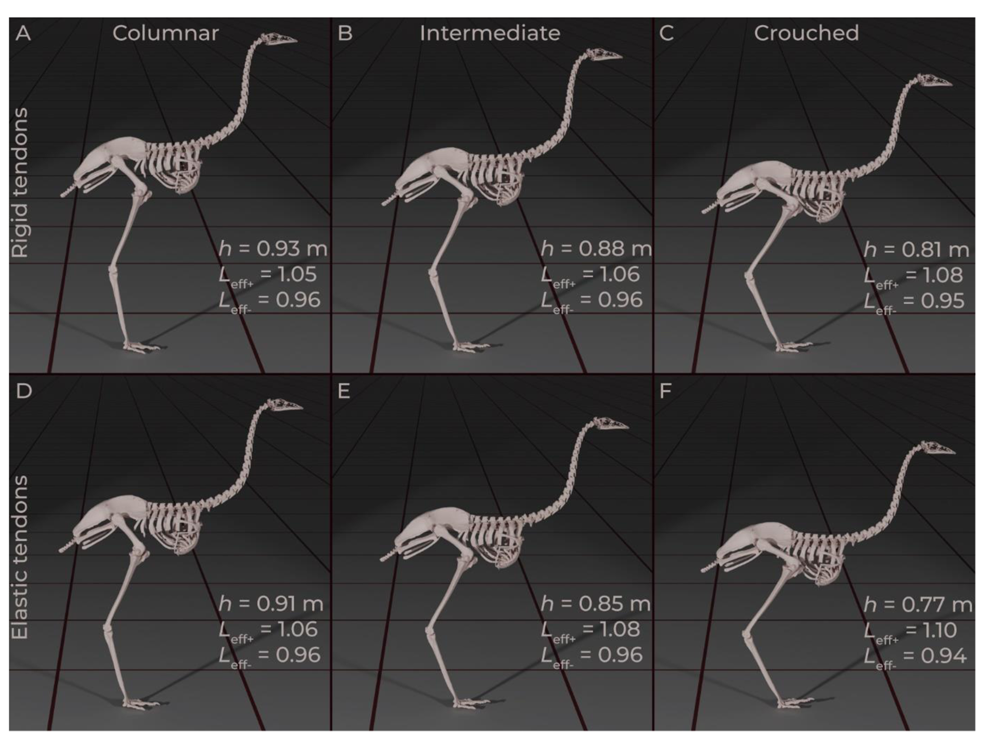
Midstance poses adopted by all the model variants when walking at 1.25 m s^-1^. From left to right are model variants tuned for columnar, intermediate, and crouched poses. The top row shows poses adopted with rigid tendons, and the bottom row with elastic tendons. Each elastic tendon variant adopted slightly more crouched postures than its rigid tendon counterpart, signified by hip height *h*. Across the walk to run transition, variations in effective leg length (*L*eff+ and *L*eff-) increased, both with increasingly crouched postures, and with elastic tendons.

Despite not having moveable toes, ground reaction forces (*GRFs*) of all model variants compared favorably to that of real emus over a large range of speeds and gaits (Figure 4, Supplementary Figures S1 and S2). Figure 4 reports both absolute speeds (*v*, in m s_-1_), and relative velocities (*v^*, dimensionless size-normalized speed, see methods). At 1.25 m s^-1^, the vertical component showed a double hump, distinctive to emus, ostriches, and bipeds in general (*13*, *26*, *49*). Irregularities in the *GRF* can be seen at higher speeds – these are related to sequential loading and unloading of neighboring contact spheres. The rigid intermediate model adopted a gait with high impact transients at running speeds (Figure S2), related to initially loading the caudal-most contact sphere (similar to a human heel-strike). Overall, our models appeared to capture the salient speed-dependent features of emu locomotion.

**Figure 4.**
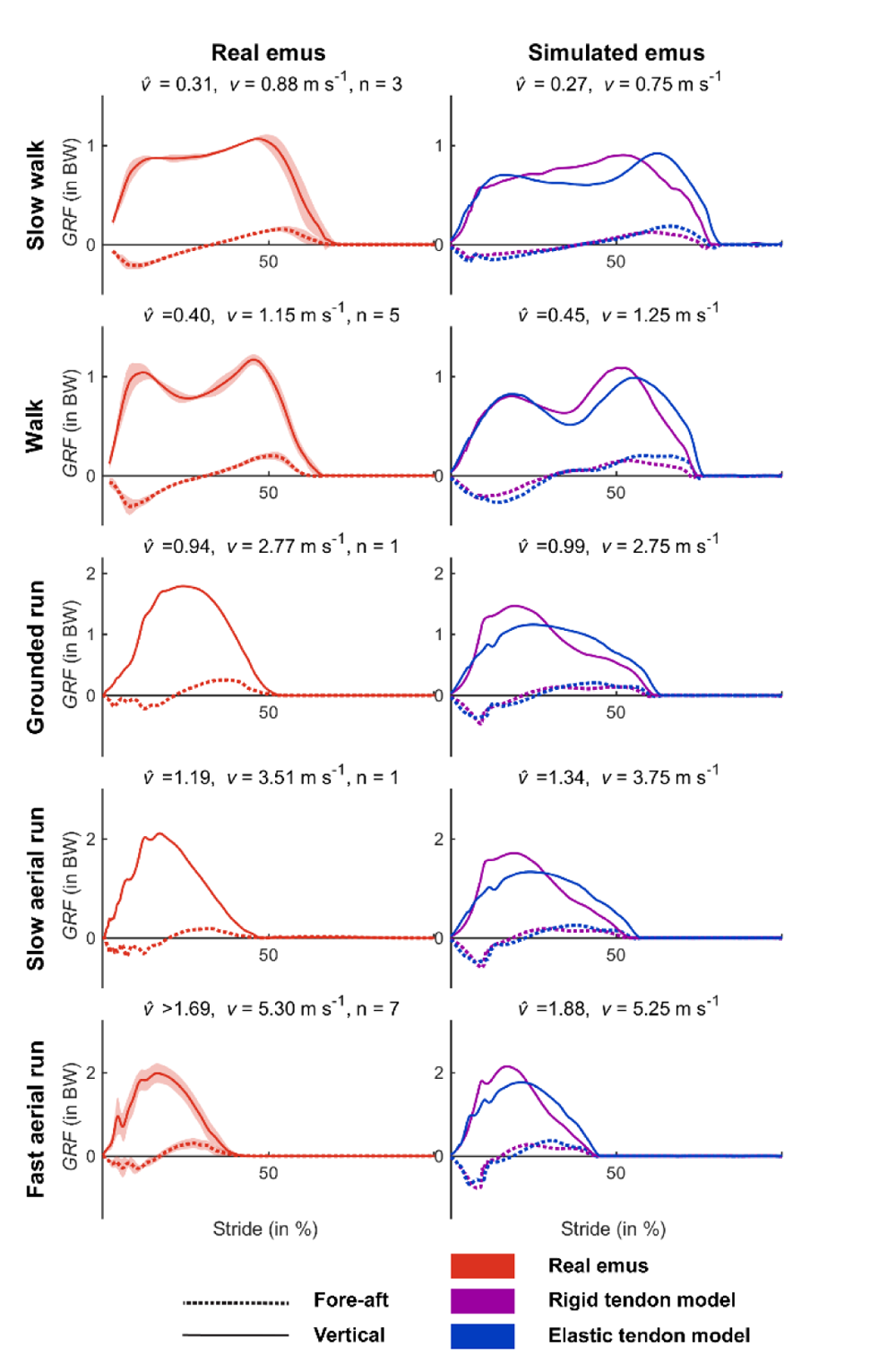
Ground reaction forces (normalized to bodyweight) of real emus (left) compared to our simulations (right) at dynamically similar speeds. We plotted the walk to run optimizations of the model tuned for crouched postures, see Supplementary Figures S1 and S2 for columnar and intermediate postures. Both model variants show the distinctive double-hump in the vertical component during walking. Two slow walk and all walk trials courtesy of J. Goetz. Grounded and slow aerial run trial, and one slow walk trial courtesy of J. Hutchinson. Fast aerial run trials courtesy of R. Main.

Duty factor (*DF*, relative ground contact time) was consistently higher in crouched model variants, than in their columnar counterparts (Figures 5 A&C, 6 A, Supplementary Figures S3 and S4). Similarly, *DF*s were higher and stride lengths were longer in the elastic tendon model variants than in the rigid tendon variants (Figure 5). Taken together, this suggests that increasingly crouched postures increase ground contact times (*DF*), and that elastic tendons have an independent effect due to their effect on postures (see “no elastic storage” sensitivity analysis).

**Figure 5.**
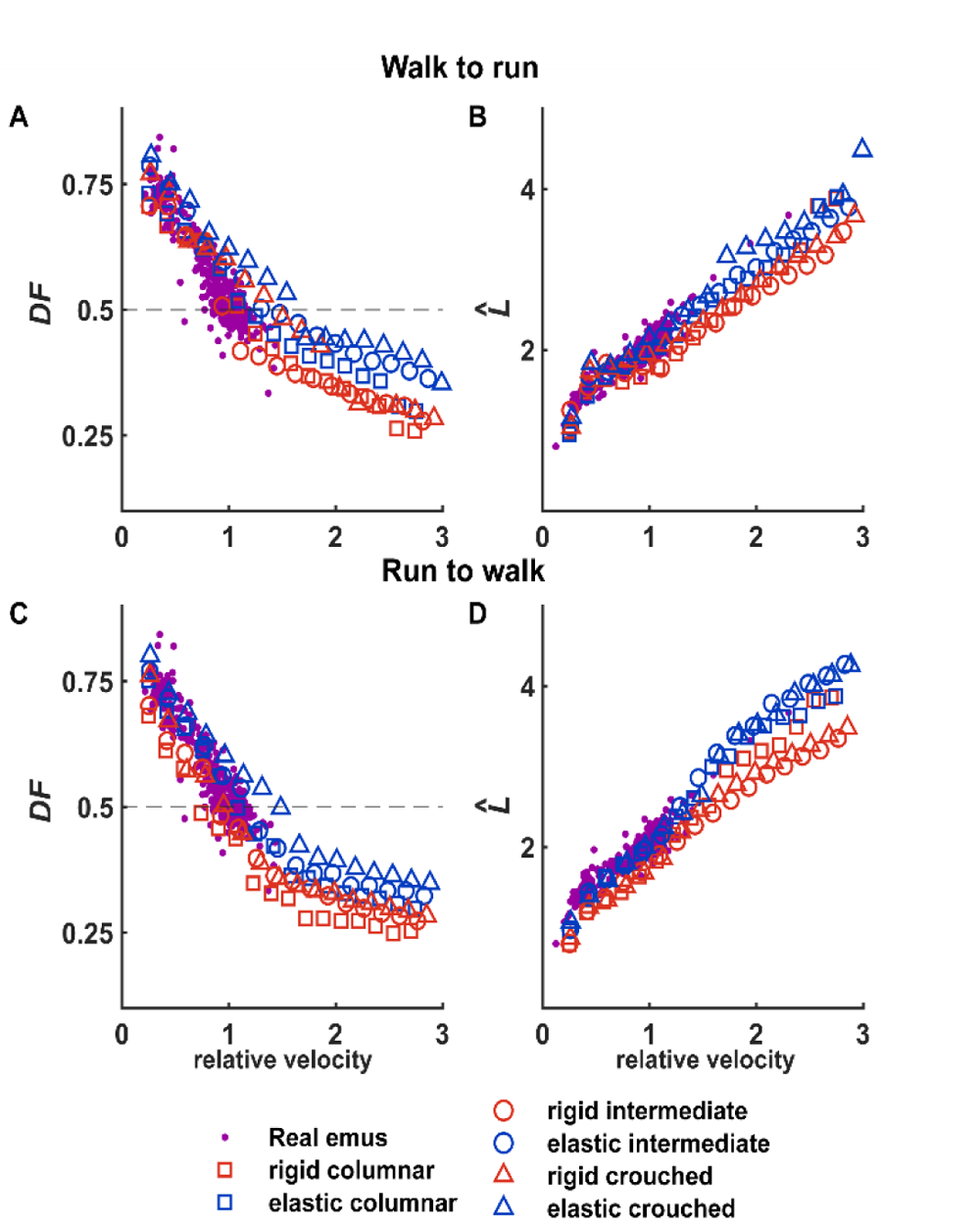
Duty factor and relative stride lengths adopted by real emus compared to our simulations. We acquired different gaits depending on whether we sequentially increased speed from walking speeds (“Walk to run”, panels A and B), or whether we sequentially decreased speeds from the maximal speed (“Run to walk”, panels C and D). Rigid tendon models (red) adopted lower duty factors and stride lengths than elastic tendon variants (blue). Columnar model variants adopted lower duty factors than crouched variants, but this did not (consistently) affect stride lengths.

### Directional hysteresis of *DF* and *L^*

We observed a hysteresis depending on whether the starting point of the sequential gait optimizations was at walking speed or maximal running speed (compare “walk to run” with “run to walk” trials, Figures 5 A&C and 6 A, Supplementary Figures S3 and S4). Thus, we acquired two gaits at each speed, which near the transition speed could be both aerial or grounded running (similar to SLIP-model predictions (*30*)). When walking (and thus high *DF*) was the starting point, the models adopted higher *DFs* across the transition speeds, and vice-versa. We plot them separately, because some model variants only adopted grounded running in the “walk to run” sequence (next section). See the Discussion for an elaboration upon this hysteresis effect.

### Gait transitions and grounded running

We defined running as a gait where the phase angle of the center of mass (*ϕ*COM, in degrees) is less than 10°. Figure 6 B shows that our models transitioned from walking to running at *v^* ∼ 0.7 – 1.1. This is slightly higher than sub-adult emus, which are reported to switch to running at *v^* ∼ 0.66 (*6*), although transition speeds in that study were determined visually. Transition speeds were similar in the fatigue-only optimizations, but were lower in the *MCOT-* only optimizations (Supplementary Figures S3 and S4, respectively).

**Figure 6.**
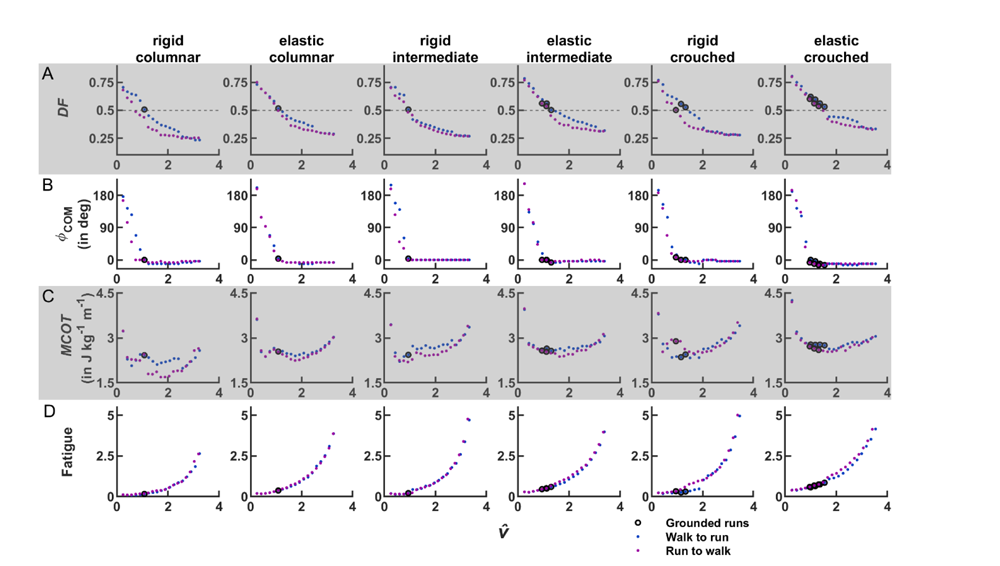
Gait metrics plotted against locomotor speed, compared between model variants. These converged optimizations used both Fatigue and *MCOT* in the cost function. Each column represents a model variant, with rows from top to bottom showing Duty factors, phase angle of the *COM*, metabolic cost of transport, and muscle fatigue. We plot non-dimensional relative velocities (*v^*). In all cases, relative velocities corresponded to a forward velocity range of 0.75 m s^-1^ – 9.75 m s^-1^, but differences in hip height affected relative velocity. Grounded running trials are indicated with black circles. The grounded runs are concentrated towards the right side of the figure, suggesting that crouched postures increase the occurrence of grounded running. Elastic tendons show an interactive effect, further increasing the occurrence of grounded running. Fatigue is normalized and scaled (see methods). Similar to figure 6, it can be seen that the models adopted different gaits depending on the optimization direction.

The elastic crouched model adopted grounded running in the *MCOT*-only optimizations (Supplementary Figure S4). However, the rigid tendon models did not display predictable gait transitions (resulting in skipping gaits or never transitioning at all), and were therefore rejected from our analysis.

Grounded running occurred when *ϕ*_COM_ was less than 10° (i.e. in-phase), combined with a *DF* of 0.5 or higher. All model variants adopted grounded running (Figure 6). In Figure 6, the occurrence of grounded runs increases from the left to right panels (black circles). This suggests that increasingly crouched postures result in grounded running being the optimal gait over a larger range of speeds. Figure 6 also demonstrates an independent effect of elastic tendons (e.g. compare the rigid crouched to the elastic crouched column): the presence of elastic tendons also increased the occurrences of grounded running. This pattern was also observable when fatigue was the singular main cost in the optimizations (Supplementary Figure S3). In the *MCOT*-only simulations, the crouched model variant adopted grounded running, while the other model variants did not. Taken together, these results suggest grounded running is increasingly advantageous with more crouched postures. The addition of elastic tendons results in more crouched mid-stance postures and larger ranges in *L*_eff_ (Figure 3), strengthening this effect (see “no elastic storage” sensitivity analysis).

Figure 6 C shows *MCOT*, which should mainly be used to compare relative differences between model variants. Because of the deliberately narrow tuning ranges of our models, our model underestimates *MCOT* of emus (*6*) (see the “wide range” sensitivity analysis and Discussion). Nevertheless, a pattern quite similar to real emus can be observed: a local optimum at low speeds (albeit at a higher speed in our simulations), then another optimum near 5 m s^-1^ (*6*). This pattern was less clear in the more crouched model variants, suggesting that these models made less use of pendular energy savings during walking. Across all simulations, minimum *MCOT* was sequentially higher with more crouched postures (Figure 6 C, Supplementary Figures S3 and S4), similar to humans (*24*). This is an important observation: given that minimal energy cost was one of the explicit optimization goals, this implies that grounded running is an energetic optimum for a crouched biped, despite having an overall higher energetic cost than columnar bipeds.

We also plot our measure of fatigue (normalized and scaled according to Equation (**2**), see methods) in Figure 6 D. It can be seen that increasingly crouched postures required higher muscle activity, resulting in higher fatigue. An analogous argumentation to the previous paragraph applies here: even though columnar tuned bipeds run with lower overall fatigue (Figure 6 D, left panels), grounded running represents an optimal solution for a crouched biped from a muscle-activity perspective.

Taken together, these results imply that grounded running is an attractive running style for a crouched biped (such as a bird), both from an energetic (*MCOT*) perspective, and from a muscle activation (fatigue) perspective. In our optimizations, relative contributions of fatigue and *MCOT* were tuned at walking speed (see methods). Fatigue increased curvilinearly with speed (Figure 6 D), whereas *MCOT* did not (Figure 6 C), resulting in an overemphasis on fatigue reduction at higher speeds. In sensitivity analysis four (below), we demonstrate that this does not affect our conclusions.

### Sensitivity analysis results

This section follows the same order as the Sensitivity Analysis subsection in the in the Methods section.

1. Our rigid tendon models could potentially store energy in the ground contacts and muscle fibers (PEE) themselves. Our “no elastic storage” model was still capable of grounded running (Supplementary Figure S5). This demonstrates that grounded running requires no elastic storage (or compliance), it is purely an effect of changes in effective leg length (*L*_eff_).
2. Our main models were deliberately tuned for narrow joint ranges, to achieve the different postures, and therefore may systematically underestimate *MCOT* (see methods). The elastic “wide range” model, was tuned for wide joint ranges to investigate this effect (Supplementary Figure S6). This model adopted grounded running over four different speeds (Supplementary Figure S6), behaving similar to the elastic crouched model (Figure 6, rightmost column). Although hip height (*h* = 0.84 m) was closer to our elastic intermediate model (Figure 3 E), the range in effective leg length was much larger (*L*_eff+_ = 1.10 and *L*_eff-_ = 0.95), similar to the elastic crouched model (Figure 3 F). This further emphasizes the significance of *L*_eff_ when interpreting grounded running. This model had higher *MCOT* over its entire range (Supplementary Figure S6), confirming that our narrow tuning ranges resulted in lower *MCOT* in the main model variants.
3. The “knee flexor” model variants investigated how the uncertain muscle function of M. femorotibialis medialis might affect gait transitions. We only simulated rigid intermediate and crouched variants (Supplementary Figure S7). The addition of the knee flexor reduced both overall fatigue and *MCOT* at higher speeds. The crouched variant still adopted grounded running, but the intermediate variant did not, which is consistent with the interpretation that grounded running is optimal in more crouched tunings.
4. Fatigue increased disproportionately to *MCOT* with increasing speeds (Figure 6 C, D). To achieve equal weighting of Fatigue and *MCOT* at 3.25 m s_-1_, relative weight of *MCOT* was increased by a factor of 2.8. Using this rescaled cost function, we ran two more optimizations with a target speed of 3.25 m s^-1^. For the crouched model, we used two solutions at 9.75 m s^-1^ as the initial guess (*DF* = 0.33 and 0.23). This still resulted in grounded gaits for the crouched model (*DF* = 0.53 and 0.52, respectively). These values for *DF* are lower than in Figure 6, but demonstrated a tendency for this model to adopt higher *DF*s. For the columnar model, we reinitialized the optimization using a 9.75 m s^-1^, and a 2.25 m s^-1^ gait (*DF* = 0.23 and 0.62). In both cases, the columnar model switched to aerial gaits (*DF* = 0.48 and 0.47, respectively). This analysis further bolsters the finding that crouched models tend towards higher *DF* gaits and grounded running, whereas columnar models do not. It also suggests, similar to the *MCOT* only optimizations, that higher weighting of *MCOT* results in lower *DF*, at the same absolute speed, which would result in the walk to run transition to occur at a lower speed.

## Discussion

Our main goal was to investigate why birds prefer grounded running at intermediate speeds. We achieved this by decoupling the effects of limb postures and tendon elastic energy storage on simulated gait transitions in the emu, using virtual experiments with a musculoskeletal model. We simulated model variants that were tuned to generate peak muscle forces at different postures (columnar, intermediate, and crouched), with both elastic and rigid tendons (Figure 3). This approach enabled us to bridge the gap between the mechanical and metabolic explanations for why birds habitually use this gait. We have shown that grounded running gaits are possible (and even optimal) in the absence of elastic tendons (Figure 6, Supplementary Figure S5). Thus, elastic energy storage in tendons, an important feature of bird locomotion, is not a requirement for grounded running. Crouched model variants adopted grounded running (and higher *DF*s in general) over a wider speed range than columnar model variants. This result was acquired regardless of whether the optimization goal was minimal *MCOT* (Figure S4), muscle fatigue (parametrized based on peak neural input) (Figure S5), or both (Figure 6). This suggests that the avian tendency towards grounded running is not paradoxical. It is an attractive running style for animals whose muscles function optimally in crouched postures, and is in agreement with well-established *MCOT*/fatigue minimization strategies. Although the evolutionary history of avian leg postures and body shapes is currently still being debated, our simulations support the interpretation that avian grounded running first evolved within non-avian theropods. We will discuss these results and their implications in turn.

### Grounded running requires changes in effective leg length, not compliance

Seminal work by McMahon et al. (*24*) related the thigh angle in running humans to leg compliance, and SLIP-models suggest that grounded running requires leg compliance (*30*). Unfortunately, such a single angle-to-compliance mapping is not generalizable to the three-segment legs of birds. More generally, compliance implies energy storage in the spring of SLIP-model, which has no clear physical interpretation. Our simulations demonstrate that energy storage is *not* a requirement for grounded running (Figure 6, Supplementary Figure S5): the unifying metric is the range in effective leg length during the stance phase.

For fixed joint excursions, the change in effective leg length will be higher at increasingly crouched postures, in absolute but especially in relative terms (Figure 3, compare A to C). This explains why our crouched models more readily lend themselves to grounded running (Figure 6 B). No storage in the abstract “leg spring” is required, and a telescopic leg actuated by a motor would likely also be capable of grounded running. However, tendon elasticity does shift the joint range at which a muscle produces maximal force (*37*), which results in more crouched midstance postures and a higher range in effective leg length (Figure 3, top versus bottom row). Thus, having very compliant or long tendons could still bias a biped with columnar optimal postures to adopt grounded running (e.g. Figure 6, elastic columnar).

### Why is grounded running in birds optimal?

Grounded running is an energetically costlier gait than aerial running (*24*, *28*), yet birds adopt it habitually at intermediate speeds (*5*, *13*, *25*, *27*). The apparent paradox, given the higher energetic costs associated with such crouched postures, can also be observed in our different model variants, since both *MCOT* and fatigue increased with more crouched postures (Figure 6 C&D, Supplementary Figures S3 and S4). However, *MCOT* and fatigue were express costs (to be minimized) during the optimizations. This suggests that grounded running gaits, and higher *DF* gaits in general, represent (locally) optimal gaits for crouched bipeds from a muscle energetics and fatigue perspective (Figures 5 A,C, 6). Crouched models converged to higher *DF* gaits than the columnar models, even if extremely fast aerial running was the initial guess (sensitivity analysis four), further supporting this interpretation. Overall, our analysis demonstrates two competing strategies for energy savings in locomotion, and that habitual grounded running in birds can be understood from that perspective.

The first energy savings strategy is to maintain as columnar a midstance posture as possible (*24*), reducing the required joint moment to resist gravity. A second strategy for energy savings is to run with higher *DF*: distributing the vertical impulse of the *GRF* over more time enables lower peak *GRF*s and lowers rates of force production in the muscles, both reducing metabolic costs (*4*, *59*, *60*). However, the second strategy also has mechanisms that increase energy cost: the prerequisite crouched postures require higher joint torques and muscle forces (*24*). The second strategy thus represents a tradeoff between minimizing peak muscle forces and rate of force development. Humans, despite being tuned for crouched postures (*38*), can avoid this tradeoff because our *COM* is situated directly above our legs, and thus a near-vertical leg posture is possible (the first strategy).

Birds cannot make use of the first strategy: in smaller birds, a vertical leg posture is impossible due to the *COM* placement in front of the hip (*36*, *40*). In ratites, *COM* placement is further backwards than in volant birds (*40*) (Figure 2 B), but a human-like posture is still impossible: our cadaveric manipulations revealed that full knee-extension in the emu is limited by as much as 42° (Supplementary Figure S8), similar to ostriches (*61*). Thus, most birds cannot save energy by adopting (human-like) extended postures, and are mechanically forced into crouched postures. Because there are now fewer negative consequences of high *DF*s, the second energy savings strategy (running with higher *DF*) can start to dominate, depending on how the tradeoff between peak muscle force and rate of force development balances out. Our simulations show that for birds, grounded running is optimal over a wider-range of speeds at increasingly crouched postural tunings (Figure 6 C,D, left to right).

A systematic evaluation of bird muscle tunings and optimal joint postures (such as (*37*) for guineafowl) has never been attempted. However, crouched poses are a feature of most extant birds, with some outliers and a tendency to adopt less-crouched postures with increasing size (*13*). Birds appear to be capable of generating forces over wide joint ranges: similar to (*22*), we have observed ratites habitually standing up from very deep squatting postures. Given that muscles are strongest at the midpoint of their range, wide active joint ranges imply somewhat crouched optimal poses for most birds. In crouched birds, the tradeoff between minimizing peak muscle force (columnar) and rate of force development (crouched) shifts towards the latter: these are the postures near which its muscles can most effectively generate the required moments (*37*), and this effect apparently dominates. This is why our crouched model variants (and the “wide range” model) adopted grounded running so frequently (Figure 6, Supplementary Figure S6).

It is important to point out that habitual postures of animals are determined by more factors than simply the posture that maximizes muscle forces (*37*, *51*, *62*). This is especially true when joint moments are low (e.g., standing or walking): passive structures, such as described for the ostrich ankle joint (*63*) and flamingo knee joint (*64*), are likely to influence habitual postures. In contrast, at higher speeds, long extensor tendons that stretch under loads lead to wider joint operating ranges (*9*, *18*), but also a more flexed midpoint (*37*) (Figure 3). Postural control also has a neural component: guineafowl adopt more crouched postures after undergoing surgery that eliminates proprioceptive feedback from their ankle extensors (*62*).

Overall, our analysis provides an elegant explanation for why it is advantageous for a bird to habitually adopt grounded running: bird muscles function optimally in relatively crouched postures, and are physically incapable of running with (near) vertical limbs. In that situation, they make the best of it by adopting high *DFs*. As such, we argue that grounded running in birds is not paradoxical: it is an attractive running style for crouched bipeds. Given the large spread in bird *COM* placements (*40*) and postures (*13*, *26*), it would be interesting to investigate whether some more columnar species adopt grounded running (e.g. the secretary bird, *Sagittarius serpentarius*).

### How did grounded running in birds evolve?

Birds are direct descendants of (non-avian) theropod dinosaurs, who relied heavily on their massive tails to power locomotion (*65–67*). Birds lost this massive tail – and resulting cranial shift in *COM* likely contributed to more crouched postures being required to stand in equilibrium (*40*). Our analysis implies that this cranial shift in *COM* was also accompanied by a strong preference for grounded running at intermediate speeds.

Given the similar morphologies, grounded running may have already occurred within the non-avian theropods. Unlike birds, non-avian dinosaurs are often thought to have adopted (relatively) columnar limb poses (*66*). Although this could suggest that they avoided grounded running, their *COM* is often reconstructed to lie in front of the hip (*40*, *68*). This situation resembles our columnar model variant, which adopted grounded running (albeit on a much narrower range of speeds than the crouched model variant, Figure 6). Fossil footprints of dinosaurs demonstrate a smooth distribution in stride lengths, which provides an independent line of evidence for non-avian theropod grounded running (*69*). Slightly problematic in this regard is that grounded running and smooth gait transitions are not always seen together (e.g. ostriches gait transitions are non-smooth (*20*), and see our Figure 6 A, elastic crouched).

Grounded running may also be more attractive at large sizes: *Tyrannosaurus rex* was observed to have adopted grounded running in predictive simulations similar to ours, although the gait only occurred at speeds where skeletal stress was higher than physiologically sustainable (*65*). Empirical data in large quadrupeds also seems to support this possibility: elephants are known to adopt running without an aerial phase (*70*, *71*).

Future simulation work on extinct taxa would be required to untangle the effects of *COM* location, leg posture, and body mass reduction on the evolutionary history of birds. Soft-tissue information is often lacking in biomechanical analyses of extinct taxa (*48*, *67*, *72*), but our simulations are encouraging in that respect: despite numerous simplifications regarding the contractile and other soft-tissue anatomy, our simulations are able to capture salient features of emu gait dynamics (Figures 4 and 5). Our results suggest that although muscular uncertainties can impact the transition speeds and maximal performance, many spatiotemporal variables (such as stride lengths) are relatively unaffected and remain similar to experimental data.

### Complexities in animal gait selection

Currently, relatively little information exists regarding how animals choose their gaits when changing speeds. Our approach has modelled changes in the desired speed as an optimization, using the current speed as a starting point. This results in a gait hysteresis – in our simulations, the walk to run transition occurs at a higher speed than the run to walk transition (Figure 6), which appears to be phenomenologically valid based on observations in the ostrich (*20*). However, we do not claim to have modelled the underlying mechanism. The gait hysteresis we found is likely an effect of using gradient-based optimizations, which may lead to minimal changes in gait dynamics when reinitializing subsequent optimizations. Put more simply, sequentially increasing or decreasing the target speed, while using each previous (slower or faster) trial as a new initial guess, likely biases the gaits to be more similar to the previous trial. This is particularly important at speeds where many different gaits are possible without a large change in *MCOT* or fatigue.

Our simulations provide an energetic mechanism that explains a preference for grounded running that is dependent on postural tuning. However, we have only modelled steady-state, level locomotion, which clearly does not encompass the full locomotor repertoire of animals (*9*, *14*, *20*). Emus habitually switch between walking and running mid-stride over a range of sub-maximal speeds (*6*), and ostriches show considerable overlap between their walking and running speeds (*20*). These and other observations suggest that animal gait selection may be situation-dependent, and indeed many factors beyond *MCOT* are likely to play a role (*11*, *12*, *14*, *31*, *62*). Grounded running has many other benefits to the organism: a large range in effective leg length (or high “leg compliance”) increases stability against perturbations (*31*), and may contribute to the capacity of birds to maintain consistent leg forces when running on uneven terrain (*14*), which is thought to be an injury prevention strategy. From this perspective, our “wide range” model shows that crouched poses may be an effect of requiring wide-active joint ranges in daily life, because of the numerous benefits it confers to the avian body plan. Indeed, these and other benefits could make grounded running attractive for bioinspired robotics (*58*, *73*).

### Limitations

In our model, we have necessarily simplified the anatomy of emus. In particular, we have modelled walking and running as pure (para)-sagittal plane movements using hinge joints, a common modelling choice (*36*, *45*, *48*). It has been shown that ratite joints display clear out-of-plane movements, particularly at the hip joint which is externally rotated throughout the stride (*74*). However, the resultant moments about the global *x* and *y*-axes (see Figure 2) have been proposed to (partially) be counteracted by passive structures (*34*), and the mediolateral component of the *GRF* does not vary systematically with speed in birds (*26*). As such, modelling these out-of-plane movements appears not to be necessary to predict overall gait dynamics, which is bolstered by the close match between our simulations and measured gait metrics in emus (Figure 4 and 5).

Our model also does not permit any movement in the metatarso-phalangeal (MTP) joints. Although the MTP joints show the largest joint excursions during the stride, this occurs predominantly during the swing phase (*27*, *74*). The *GRF* progressions of our models (Figure 4) demonstrate that omitting the MTP joint does not prevent realistic foot-ground interactions, and thus overall stance-phase dynamics, which seem most relevant to grounded running.

Several adaptive features linked to low *MCOT* in ratites and birds are not included in our model: ankle ligaments generate moments that lead to two stable joint poses in ratites (*63*), and the digital flexors store large amounts of energy during the stride (*9*, *18*, *34*, *58*). Emu hindlimb muscles are also metabolically and structurally highly-specialized for sustained high-speed running (at low energy cost) (*19*, *33*, *57*), which is not represented in our model (Figure 2). In fact, lumped parameter Hill-type muscles may overestimate fiber length changes (*75*), which in our methodology would lead to weaker muscles when keeping muscle mass constant (equation (**1**)).

It is thus reasonable to assume that our model would underestimate the locomotor performance of emus by omitting these features. It may then seem surprising that most of our model variants have lower *MCOT* than reported for emus (*6*). We remind the reader that we tuned our model variants for narrow joint ranges, to deliberately bias it towards specific postures – using wide joint ranges would prevent us from performing the main analysis (see Muscle Contractile Parameters, in the methods). At a fixed muscle mass, these narrow joint ranges result in relatively short fibers (low *L*O), which per (equation (**1**)) leads to strong muscles (high *F*_max_). An effect that is strengthened because we mapped individual muscle masses onto functional groups in the sagittal plane. As such, the values of *MCOT* in Figure 6 C are underestimates. Our “wide range” model from the sensitivity analysis shows that a wider tuning range results in higher *MCOT*s (Supplementary Figure S6). To accurately model all aspects of ratite locomotor performance simultaneously (wide active joint ranges, high top speed, low *MCOT*), it may be necessary to model all muscles individually, or even split up each muscle further (e.g. (*37*, *49*)) – substantially complicating the control procedure and increasing computational time. Muscle power output in mice decreases under repetitive contractions (*76*), and similar experiments have not been performed on ratite muscles. As such, we currently lack sufficient anatomical and physiological information to simultaneously capture all aspects of ratite muscle functioning using computational models.

Lastly, as is often the case with simulation studies, it is impossible to guarantee global optimality of the presented gaits, demonstrated by the differences between our “walk to run” and “run to walk” optimizations discussed previously. We have taken numerous steps to investigate the robustness of our claims, including systematically searching for alternative walking solutions, approaching the gait transition speeds from two directions, and investigating the effect of different cost functions. A recurring pattern in all of these investigations was that models tuned for crouched postures adopted higher *DF*s.

## Conclusions

We have presented a selection of emu model variants that enabled us to isolate the effect of different aspects of their musculoskeletal design on overall gait dynamics. By simulating grounded running both with and without elastic tendons, we demonstrated that grounded running does not require elastic storage, although it is facilitated by it. Mechanically, grounded running gaits require larger variations in effective leg length during the stance phase than aerial running – this is easiest to achieve with a crouched posture. We demonstrated this by simulating different emu model variants that were tuned to generate the highest muscle force on a gradient from columnar to crouched postures. Our model variants converged to grounded running over the widest range of speeds when tuned for a crouched posture. More generally, we found that models that were strongest in crouched postures, adopted crouched running gaits with higher duty factors than columnar models. Thus, while columnar postures enable lower joint torques (and thus lower muscle forces), columnar running is only advantageous when the muscles function optimally in these postures. Humans are an exception in this regard, because we can stand in equilibrium when we fully extend all of our joints. In birds, a fully extended posture is impossible due to the forward placement of the center of mass. Integrating these findings, we reject the existence of a grounded running paradox in birds. We conclude that birds adopt grounded running because it is the most energetically advantageous gait that is available to them.

## Methods

### Overview

We have constructed a generalizable (i.e., not subject-specific) 3D musculoskeletal model of the emu for use in predictive physics simulations. The model incorporates all the unique mono-and bi-articular (sagittal plane) muscle functions of emus (*19*, *33*), while combining muscles with identical functions into functional groups. We systematically varied the muscle contractile parameters so peak muscle forces in the model occurred in three different postures (combinations of joint angles): columnar (albeit less extended than humans), intermediate, and crouched (Figure 3). For each of these three postural variants, we considered the effects of both rigid and elastic tendons, resulting in six model variants in our main analysis.

Using optimal control theory, we optimized gaits for the model variants, across their entire speed ranges. Importantly, our simulations were acquired without using movement data from emus as an *input.* Systematic differences between model gaits thus reflected differences in musculoskeletal design (posture and/or tendon elasticity) of the models. To validate simulator *outputs*, we compared them to published gait data of real emus (ground reaction forces, stride lengths, *DF*).

Because optimizations can be sensitive to initial conditions, we simulated gait transitions in two directions. “Walk to run” – this modality started from walking speed, and we increased the target speed for subsequent trials by 0.5 m s^-1^ until we found the model variant’s top speed. See Video S1. “Run to walk” – this modality started at top speed, and we decreased the target speed for each trial in steps of 0.5 m s^-1^. Each trial used the previous trial from the sequence as the initial guess for the optimization at the new target speed.

In the simulated gait transitions acquired in this way, we compared how posture and tendon elasticity affected the occurrence of grounded running, and corresponding metrics of *MCOT* and muscle fatigue. We also performed extensive sensitivity analyses to investigate the robustness of our claims to implementation choices. This included the inclusion of alternative cost functions (in the optimizer), and several attempts to acquire aerial gaits at speeds where grounded gaits were optimal, and vice versa. In total, we have analyzed the results of over 650 optimizations.

We provide code, all model variants, and simulator outputs as supplementary data, and have also uploaded them to an open repository (see Data Availability).

### Inertial parameters

We acquired a CT-based 3D model of an emu skeleton from (*40*). We imported the 3D models into Blender 3.0 (blendernation.org, open-source 3D modelling software). To account for the posture it was scanned in, the skeleton was reposed into a neutral position, and the torso was slightly symmetrized (Figure 2 A, top). We then applied a validated convex hulling procedure (in Blender) to estimate the inertial parameters directly from the skeleton (*40*). The basic procedure is as follows:

1. generate minimal (3D) convex hulls around distinct body segments as defined in (*40*).
2. scale the hulls using segment-specific scale factors validated in birds (*40*), so that it represents the skin outline; the skeleton was not perfectly symmetric – we generated symmetric hulls by mirroring the right-side to the left-side, while accounting for the change in volume due to this extra step.
3. multiply the volumes and volumetric inertia tensors by segment density to obtain the mass properties.

We used the average (isometric) bird convex hull expansions from (*40*). Expanding the torso as a single segment redistributes an inappropriate amount of volume around the ribcage. After computing the total torso volume as predicted by their equations, we split the unexpanded torso hull into a pelvic-portion, and a ribcage-portion. We only expanded the ribcage-portion enough to ensure the symmetrized hull enveloped the skeleton. The rest of the torso volume was accounted for by expanding the pelvic-portion. Splitting up the torso segment in this way results in a torso skin outline that more closely resembles the CT-based skin outlines (*40*), where the pelvis is much wider than the ribcage. To more closely match the proximal concentration of mass in the thigh and shank segments of emus, the corresponding hulls were only expanded on the proximal ends until the target volume was achieved. The final hulls are shown in Figure 2 A (bottom). All hulls were assigned a density of 1000 kg m^-3^, except for the head, neck and torso hulls, which were assigned a density of 888 kg m^-3^ (*68*), and the calculated body segment parameters are presented in Table 1. Estimated body mass was 37.8 kg, which was within 1% of the mass measured using scales (38.1 kg) for the same individual (*40*).

### Muscle masses

We compiled a dataset of bodyweight-normalized muscle masses of the average adult emu hindlimb, based on published data. Lamas et al. (*33*) reported two large datasets, ranging from hatchlings to adults, which we combined with an adult measurement from Goetz et al. (*49*). We adopt the standardized nomenclature of the Handbook of Avian Anatomy (*77*), but some of the data followed nomenclature of Patak and Baldwin (*19*). These nomenclatural inconsistencies are also present in ostrich myology, briefly discussed in (*51*). We converted all the data to a single naming convention, and excluded uncontroversially juvenile specimens (based on muscle mass, see supplementary texts). This muscle mass dataset is based on 28 specimens (body mass range: 15.6 – 51.7 kg), with sample sizes varying per muscle (Table 2). The median sample size was n = 19 (for nine muscles), the lowest was n = 9 (ISF, which is a diminutive muscle that is challenging to dissect, see (*33*)), and the highest was n = 28 (for FL, EDL, FMTL, GM, GL, and GIM). Hindlimb muscle mass of emus (raised in captivity, excluding juveniles) in this dataset is on average 14.9% of total body mass (per limb), nearly identical to the 15% reported in Lamas et al. based on a subset of their own data (*33*). The highest percentage (16% body mass per limb) was found in a specimen from their UK dataset (at a body mass of 42 kg). Patak and Baldwin (*57*) report ankle-extensor muscle masses for wild emus, which on average are 7% heavier than the captive emu dataset presented here.

The 34 muscles listed in Table 2 have considerable functional specializations due to their varying architecture (*19*, *33*, *57*). Our study design relies on biasing the model towards specific poses, and representing the real functional complexity of the muscular system is counter-productive to this goal. Furthermore, our model only incorporates sagittal plane movements, because medio-lateral forces do not vary systematically with speed in birds (*26*). Therefore, we mapped the 34 muscles of the emu hindlimb onto five mono-articular, and four bi-articular functional groups, which we consider to capture the major muscle functions in the sagittal plane. These functional groups, body mass-normalized muscle masses, and the anatomical muscles they are based on, are presented in Table 3. Figure 2B shows the corresponding muscle lines of action, which were constructed using comparisons with existing ratite models (*49*, *51*) and a 3D surface scan of a *Rhea* plastinate with the muscles preserved in situ.

When a unique mapping was not appropriate, muscle masses were distributed evenly over multiple functional groups (see supplemental texts). Table 3 includes a sixth mono-articular muscle (knee flexion by M. femorotibialis medialis (*21*, *33*, *51*)). Its topographical arrangement suggests that it is predominantly a knee-adductor (or stabilizer), but some authors have also classified it as a (weak) knee-flexor (*33*, *51*). Given this uncertainty in the literature, we have added its muscle mass to the “Hip extensor, knee flexor” functional group for all the results of the main analysis. We investigated its effect as a mono-articular flexor in the sensitivity analysis.

### Muscle contractile parameters

Our goal was to assess how habitual limb postures affect locomotor dynamics. We achieved this by generating model variants with contractile parameters of the nine muscle-tendon units tuned for different postures. In animals, muscle fibers operate over relatively narrow length ranges in-vivo, and cannot actively generate large forces outside of these ranges (*10*, *39*). Optimal lengths of the muscle fibers, combined with dynamic interplay with the tendons, are highly determinant of habitual limb poses and joint ranges (*10*, *37*, *39*). Similar to previous animal simulation studies (*48*, *50*, *52*, *65*), we tuned our model using specific joint ranges (and thus postures) by using the length changes of the muscle in different postures to determine fiber and tendon lengths (explained in detail below).

We chose three limb postures to (see Figure 3, Results section, and Video S2): columnar (relatively extended joints), intermediate, and crouched (flexed joints). These postures were not intended to match emu joint angles, but represented a suitably wide postural gradient to allow us to investigate the effect of posture on gait selection. Cadaveric manipulation of several ratites (*Dromaius, Casuarius sp.* and *Rhea sp.*) revealed wide passive joint flexion/extension ranges (approximately 85° hip, 115° knee, and 165° ankle, with the last 35° of ankle flexion being resisted by passive forces). Using these cadaveric ranges as tuning ranges would be incompatible with a columnar tuning (if joint hyperextension is not allowed), because muscles are strongest at the approximate midpoint of their range. To be able to generate a columnar variant at all, it is necessary to choose relatively narrow joint tuning ranges. However, if joint ranges are too narrow, fiber lengths take on unrealistically low values (see below). After pilot testing, we found joint tuning ranges of 45° (hip), 65° (knee), and 70° (ankle) to represent a reasonable compromise (slightly more than half the unresisted passive joint ranges).

Compared to joint excursions in walking and running ratites (*27*, *49*, *63*, *74*), our joint tuning ranges are twofold wider at the hip (but similar to (*27*)), and slightly narrower (∼15°) at the ankle. This was a necessary departure from bird behavior to enable a systematic comparison of the effect of habitual posture on locomotion. To investigate the effect of this, we considered a model tuned for much wider active joint ranges as a sensitivity analysis (the “wide range” model variant).

We modelled the muscles using three-element Hill-type muscles, incorporating first-order activation dynamics (*78*, *79*). A muscle fiber is represented by a contractile element (CE) and a non-linear parallel elastic element (PEE) and generates the highest isometric force at its optimal fiber length (*L*_O_, in m). For the elastic tendon variants, tendons are represented by a non-linear series elastic element (SEE), with a corresponding tendon resting length (*L*_T_, in m). The fiber and tendon are placed in series to represent a whole muscle.

We set *L*_O_ of each muscle fiber so that it extends between 0.5 – 1.5 when the joints are moved over the tuning ranges. For the columnar model variant, we set the midpoint of these ranges at 22.5° (hip), -47.5° (knee), and 40° (ankle). *L*_T_ was determined by subtracting *L*_O_ from the total muscle length at these midpoints. We biased the model towards intermediate and crouched postures by shifting the midpoints by 10° and 20°, respectively, and recalculating *L*_O_ and *L*_T_. The tuning ranges were only used to construct the models, during gait optimizations the models were free to adopt a much wider range of joint angles. However, these tuning ranges achieved systematic differences in posture between conditions, which was our goal.

Maximum isometric force (*F*_max_, in N) was dependent on the muscle’s mass (*m*_muscle_, in kg) and *L*_O_ using equation (**1**):

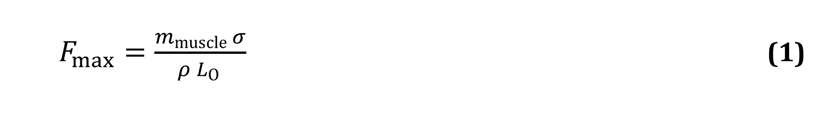

Here, *σ* is the specific tension of muscle tissue (0.3 MPa used here, (*48*, *50*, *65*)), and *ρ* is the density of muscle tissue (1060 kg m^-3^). We do not explicitly model the pennation angle, because the functional muscle groups in our model represent the combined effect of several muscles.

Given that we prescribed *m*_muscle_ for each muscle, equation (**1**) shows that *F*_max_ is inversely proportional to *L*_O_: a muscle with short fibers has a larger maximal force, and vice versa. Our deliberate choice for relatively narrow joint ranges therefore likely increases *F*_max_, causing a systematic underestimation of *MCOT*. We investigate this effect in the sensitivity analysis (“wide range” model variant).

For the elastic tendon model variants, tendon strain at *F*_max_ was set at 4%. Similar to (*37*, *52*), *L*_T_ was kept constant between the rigid and elastic tendon variants to directly compare the effect of tendon elasticity.

### Model construction

We treat the head, neck, forelimbs, and torso as a single rigid body (Table 1). The model has bilaterally symmetric rigid bodies representing the thigh, shank, and foot bodies, resulting in three-segment legs. The foot body is made up of the midfoot (tarsometatarsus) and the toes (phalanges), which cannot move independently from each other – they are fixed in a rigid L-shape (Table 1, Figure 2). Joint locations (Figure 2B, purple circles) were determined by manually fitting geometric primitives to the articular surfaces in Blender.

All the three-dimensional data (inertial properties, joint locations, muscle paths, contact sphere locations) were computed in Blender, and then converted for simulations in OpenSim 4.4, open-source software for biomechanical analysis (*80*). We constructed the model using the OpenSim API in Matlab R2019b (Mathworks, Natick). We constrained the “head, neck, forelimbs, and torso” body to the sagittal plane (three degrees of freedom), all other joints were hinge joints, leading to nine degrees of freedom in total.

The model has 9 controllable muscles per hindlimb. We used the muscle model described by De Groote et al. (*46*) (implemented in OpenSim as “DeGrooteFregly2016Muscle” (*54*)). Parameters that were kept constant for each muscle throughout our experiment are: maximal contraction velocity (14 *L*_O_ s^-1^, based on turkeys (*81*)), activation time constant (15 ms), deactivation time constant (50 ms), PEE strain at *F*max (60%).

The model has 10 ground contact spheres per foot (0.015 m radius). Contact forces were modelled using a smoothed Herz-Hunt-Crossley model, as described in (*42*) (implemented in OpenSim as SmoothSphereHalfSpaceForce (*54*)). Plain strain modulus was set to 2.5 MPa, dissipation to 0.2 s m^-1^. Static and dynamic friction coefficients were set to 0.4, and viscous friction was set to 0.1 to prevent slippage at higher speeds. The “herz_smoothing” parameter was set to 600.

Model-specification files (in .osim format) of all our model variants can be found in the supplementary materials and on the SimTK Project page (see data availability statement), which can be interactively posed and examined in OpenSim 4.4.

### Optimizations

In our model, the only controllable inputs are the open loop neural inputs (excitations), one for each muscle, with no model state feedback. Excitation results in muscle activation (via first-order activation dynamics). Each muscle’s force production depends on its instantaneous fiber length and velocity, and its activation. The time dependent neural inputs of all the muscles together must result in periodic locomotion. Finding appropriate neural signals for periodic gait was traditionally extremely computationally costly (*48*, *50*). These neural signals are commonly found using optimal control: the gait of the model can be optimized by assuming that animals select their gaits by minimizing one or several objectives (*11*, *41*, *45*, *48*, *54*).

Optimizations that include *MCOT* as the only (main) objective often do not produce realistic motions (*44*, *45*), because they do not equally distribute loads over all the available muscles. By minimizing the neural input to the muscles raised to a power, muscle activity is distributed more equally across the muscles (irrespective of size), often resulting in more realistic gaits (*45*). This latter cost is thought to represent fatigue sensing in the muscles (*45*). While there is evidence that in certain situations, fatigue dominates over *MCOT* in gait selection (*11*), there is currently no consensus on their individual contributions in animals. Therefore, we used three different objectives for our optimizations: 1) Fatigue and *MCOT*; 2) only fatigue; 3) only *MCOT*. Our main analysis focuses on the combined Fatigue and *MCOT* optimizations.

Recently, direct collocation has gained popularity as a control approach with greatly reduced optimization times (*45*, *46*, *41*, *54*). Direct collocation differs from more traditional simulation workflows because trials are not acquired through forward integration over time. Instead, all the time points of the simulation are optimized simultaneously, and system dynamics are enforced as equality constraints that can initially be ignored during the optimization. We will refer to our converged optimizations (i.e. gait trials) as simulations (*39*, *41*, *45*). We implemented direct collocation using the optimization suite Moco (in OpenSim) to acquire biologically realistic gaits for our model (*54*). We set convergence and constraint tolerances in Moco to 10^-3^ and 10^-4^, respectively.

### Cost function

We implemented a multi-term objective function, similar to (*41*):

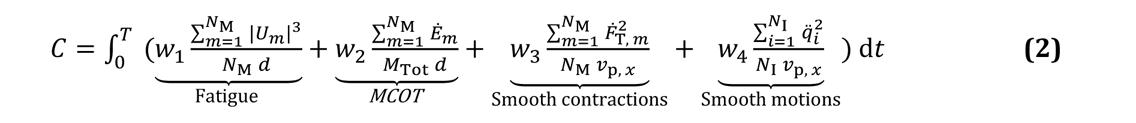

*w*_1_ – *w*_4_ = weights for scaling the four terms in the cost function

*T* = final time point (in s) of the simulation

*N*_M_ = number of muscles

*N*_I_ = number of coordinates (degrees of freedom)

*U_m_* = control input, or excitation (bounded between 0.0001 and 1), of muscle *m*

*d* = distance traversed (in m) in *x*-direction by the pelvis, between *t* = 0 and *t* = *T*

*v*_p, *x*_ = average velocity (in m s_-1_) in *x*-direction of the pelvis

*M*_Tot_ = total body mass (in kg)

*Ė_m_* = total metabolic rate (in W) of muscle *m*

*Ḟ*_T, *m*_ = time-derivative of the normalized tendon force of muscle *m*

*q_i_* = second time-derivative of coordinate *i*

Weight *w*_1_ was kept constant between all model variants. Weight *w*_2_ was selected so the contributions of fatigue and *MCOT* were approximately equal at 1.25 m s_-1_. The smoothness criteria *w*_3_ and *w*_4_ are typically used to improved numerical conditioning (*41*). They also serve to limit tendon oscillations (when using elastic tendons) and jittery gaits. We scaled *w*_3_ and *w*_4_ so that their respective contributions to *C* were small (<10% of the muscle fatigue at 1.25 m s^-1^). All weights were kept constant for all the speeds considered. *w*_3_ was zero in the rigid tendon simulations. The velocity terms multiplied by *w*_3_ and *w*_4_ ensure their relative contributions are somewhat constant irrespective of the speed, but a systematic treatment of cost-function scaling goes beyond the scope of the current study. To account for this, we also ran the optimizations without *MCOT* and without fatigue included (*w*_2_ and *w*_1_ set to zero, respectively). We also include sensitivity analyses where we rescaled *w*_1_ and *w*2 at 3.25 m s^-1^ (grounded running speed).

*MCOT* was estimated using the phenomenological model proposed by Bhargava et al. (*82*), assuming a 50% distribution in slow and fast muscle fibers. We set the basal metabolic rate to 0.92 W kg^-1^, the mean of all emu measurements reported in (*83*), assuming 20.1 J of energy released per ml^-1^ of oxygen consumption (*3*).

Each half-stride was discretized into 101 time points. This has the desirable effect of automatically decreasing the simulation time steps at faster target speeds, because the stride periods become progressively shorter.

### Constraints

We enforced target average speed in *x*-direction, and we also enforced periodicity and bilateral cross-symmetry of coordinates and muscle excitations. This results in half strides where the initial states and controls of the right side match the final states and controls of the left side, and vice-versa. We transformed these half strides into full strides by placing a bilaterally mirrored trajectory in sequence with the original. For an arbitrary state *S* that has a left and right variant, the constraints to acquire the half strides can be written as:

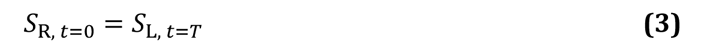

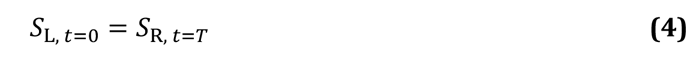

We further placed lower and upper bounds (*b*_L_ and *b*_U_, respectively) on all state variables defining position (*q*) and velocity (*q*):

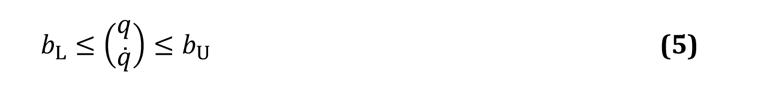

Apart from the constraint on pelvis angle, these bounds were wide enough to remain inactive in all of the converged solutions, but served to reduce the size of the search space. These bounds do not represent mechanical constraints on the model – no constraint forces would occur if the bounds are reached. Instead, these bounds should be interpreted as controller limits on acceptable joint poses and speeds.

### Gait generation

We first generated optimal walking gaits at a target speed of 1.25 m s^-1^ (close to preferred walking speed of emus (*6*)), using two initial guesses for each model iteration: 1) a dynamically *consistent*, static balance simulation; 2) a dynamically *inconsistent*, “quasi-random” guess (*41*), where the model floated forward at the target speed with swinging limbs. Both initial guesses sometimes resulted in non-optimal gaits (e.g., hopping, skipping, marching). Some examples of these are provided as a supplementary video. These local optima were avoided somewhat by initially placing tight bounds on the coordinate speeds and *T* – these bounds were lifted once a good walking solution was found. We also implemented a procedure similar to “gait morphing” (*48*) – reinitializing the optimizer with a (portion of a) converged solution as a new initial guess.

For each of the two walking solutions found per model, we further investigated whether enforcing stride lengths between 0.75 – 1.75 m (in steps of 0.25 m) would lead to better solutions (because we have multi-objective cost function). We also cross-checked solutions for the different model iterations if we suspected a local optimum. Overall, we generated at least 12 viable solutions per main model iteration, before selecting the solution with lowest cost.

We then sequentially increased the target speed in steps of 0.5 m s^-1^, each time using the previous solution as a “hot start”, until we reached a speed at which the optimizer would no longer converge. We refer to this sequence of optimizations as the “walk to run” modality (see Video S1). In these optimizations, the average velocity of the initial guess was always 0.5 m s^-1^ lower than the target velocity, which could potentially induce a bias in the gait transition speeds. To eliminate this bias, we repeated the procedure in reverse: using the fastest converged speed (always an aerial running gait) as an initial guess, we sequentially reduced the target speed until 0.75 m s^-1^. This sequence will be referred to as the “run to walk” modality.

### Sensitivity analyses

We performed extensive sensitivity analyses, in the form of extra model variants and alternative optimization approaches, which we list here.

1. Our rigid tendon models still had the potential for some elastic storage in the PEE and the smooth ground contacts. To eliminate the possibility that any elastic storage was required for grounded running, we constructed a “no elastic storage” model variant. This was a rigid crouched model variant, without a PEE and much stiffer (non-smooth) ground contacts (HuntCrossleyForce, as implemented in OpenSim (*84*)). Plain strain modulus was set to 12.5 MPa (five times higher than the base model variants). All other parameters were kept the same.
2. Our main analysis depended on deliberately tuning the model variants for narrower joint ranges than in real life (see the Muscle Contractile Parameters section). To explore beyond this experimental paradigm, we constructed a “wide range” model variant. This model was tuned with much wider joint ranges, approximating the unresisted passive joint range of emus determined through cadaveric manipulation. This model was tuned assuming 85° hip, 115° knee, and 130° ankle ranges (with midpoints at 42.5°, -82.5° and 70°, respectively).
3. The function of M. femorotibialis medialis is currently poorly understood, so our main model variants do not include its potential effect as a mono-articular knee-flexor (KF). To investigate its effect during gait, we constructed “knee flexor” model variants: intermediate and crouched model variants (with rigid tendons) with a KF muscle.
4. We initially scaled the relative weights of fatigue and *MCOT* in the cost function (*w*_1_ and *w*_2_ in equation (**3**), respectively) at a speed of 1.25 m s^-1^. However, the cost of fatigue increased more substantially at higher speeds than *MCOT*. We investigated how reweighting the cost function at 3.25 m s^-1^ (grounded running speed for the elastic columnar and crouched models) would affect the gait optima.

Unless stated otherwise, we used a walking solution of one of the main model variants that most resembled it to generate a new walking solution, before traversing the walk-to-run transition as described above.

### Computation of gait metrics

We compared gaits optimized for our model with published data on emus and ostriches (*17*, *25*– *27*, *34*, *49*). To aid these comparisons, we computed non-dimensional gait parameters as follows: we defined *h* (in m) as the average hip height during walking. We computed *h* at a forward velocity of 1.25 m s^-1^ in our simulations, but some of the studies we compared to did not report the velocity at which they determined it, or computed it for standing trials. Stride length *L* (in m) was defined as the distance between successive footfalls of the same foot. We computed relative stride length *L^* by normalizing stride length to *h*. Relative velocity *v^* was obtained by dividing forward velocity *v* (in m s^-1^) by (*g h*)^0.5^, where *g* is the gravitational acceleration constant (in m s^-2^). The quantity (*g h*)^0.5^ is also the upper limit on walking for an inverted pendulum. *v^* = 1 thus represents a theoretical upper limit on walking gaits, although grounded running at *v^* > 1 is possible in the SLIP-model (*30*) and is regularly adopted by birds (*26*). The Froude number *Fr* for pendular walking is equivalent to *v^*^2^ (*1*). Because we only focus on steady-state, forward locomotion, we will use the terms velocity and speed interchangeably.

Ground reaction force (*GRF*) was reported as a fraction of body weight. Duty factor (*DF*, fraction of the stride that one or both feet are in contact with the ground), was determined based on the presence of intersections between contact spheres and the ground.

To compare stance phase dynamics between model variants, we defined effective leg length *L*_eff_: the average distance (in m) between the center of pressure and the hip joint during the stance phase. To focus our comparison on the walk to run transition, we computed the average over a speed range of 1.25 – 3.25 m s^-1^. We report maximum and minimum values, normalized to the average (*L*_eff+_ and *L*_eff-_, respectively).

Data from (*25*, *27*, *34*) were digitized using PlotDigitizer (plotdigitizer.com). To convert the measured running speeds from Rubenson et al. (*34*) to *v^*, we used *h* = 1.09 m. This value was acquired by scaling *h* of the ostrich reported in (*25*) (1.19 m), assuming *h M*_Tot_^0.33^. Force-plate and kinematic data of emus were kindly shared with us by J. Hutchinson (from (*26*, *85*)), J. Goetz (from (*49*)) and R. Main (from (*17*)). We processed these to generate *GRF* traces of emus at a variety of speeds (see supplementary text for details), and the data from (*49*) also provided extra data points for *DF* and *L^*.

The phase angle of the *COM* oscillations (*ϕ*_COM_, in degrees) was calculated using a method similar to (*5*): we split up total *COM* energy into contributions due to horizontal and vertical velocity fluctuations of the *COM*, and summed the vertical fluctuations with the gravitational potential energy of the *COM.* We computed the cross-correlation between these two contributors to total *COM* energy. *ϕ*_COM_ corresponded to the time shift at which the cross-correlation was maximal. We do not report the absolute phase angles, but define *ϕ*_COM_ with respect to the vertical fluctuations (e.g., if *ϕ*_COM_ = 90°, the horizontal peak comes a quarter period after the vertical peak). For the purposes of our analysis, we operationalize grounded running as a gait that where *ϕ*_COM_ is less than 10° combined with a *DF* that is 0.5 or higher.

## Data availability

Models, simulator outputs, and code are freely available on our SimTK project page: https://simtk.org/projects/emily_project (which stands for Emu Model for Investigating Locomotor dYnamics). These data have also been provided as electronic supplementary data to this manuscript. Supplementary texts and figures are provided as two separate documents.

We have also produced a video abstract, and three supplementary videos: Video S1 – Demonstration of the gait generation procedure

Video S2 – Comparison of walking gaits adopted by the models

Video S3 – Examples local optima and other rejected gaits

## Acknowledgements

We thank Russel Main, Jessica Goetz, John Hutchinson and Luis Lamas for sharing their emu data, and assistance during data processing. We thank Monica Daley, Stephen Gatesy, Peter Bishop, Annick Abourachid, and Annette Koenders for helpful advice and discussions. The authors are very appreciative of discussions with Nicholas Bianco, Ross Miller, Aaron Fox, Antonie van den Bogert, Bryan Umberger and Antoine Falisse on the Moco forums, which were instrumental in shaping the methodology implemented here. We also thank Thomas Geijtenbeek, William Sellers, Anne Koelewijn, Koen Lemaire, Florian Muijres and Johan van Leeuwen for discussions on gait simulations and mechanics in general. Lina Walen is thanked for providing specimen access. Lastly, the authors thank Victor Duurland for providing permission to photograph his emus, and Meret Spithoven and Matt Dempsey for assistance during data collection.

---

**Figure S1.**
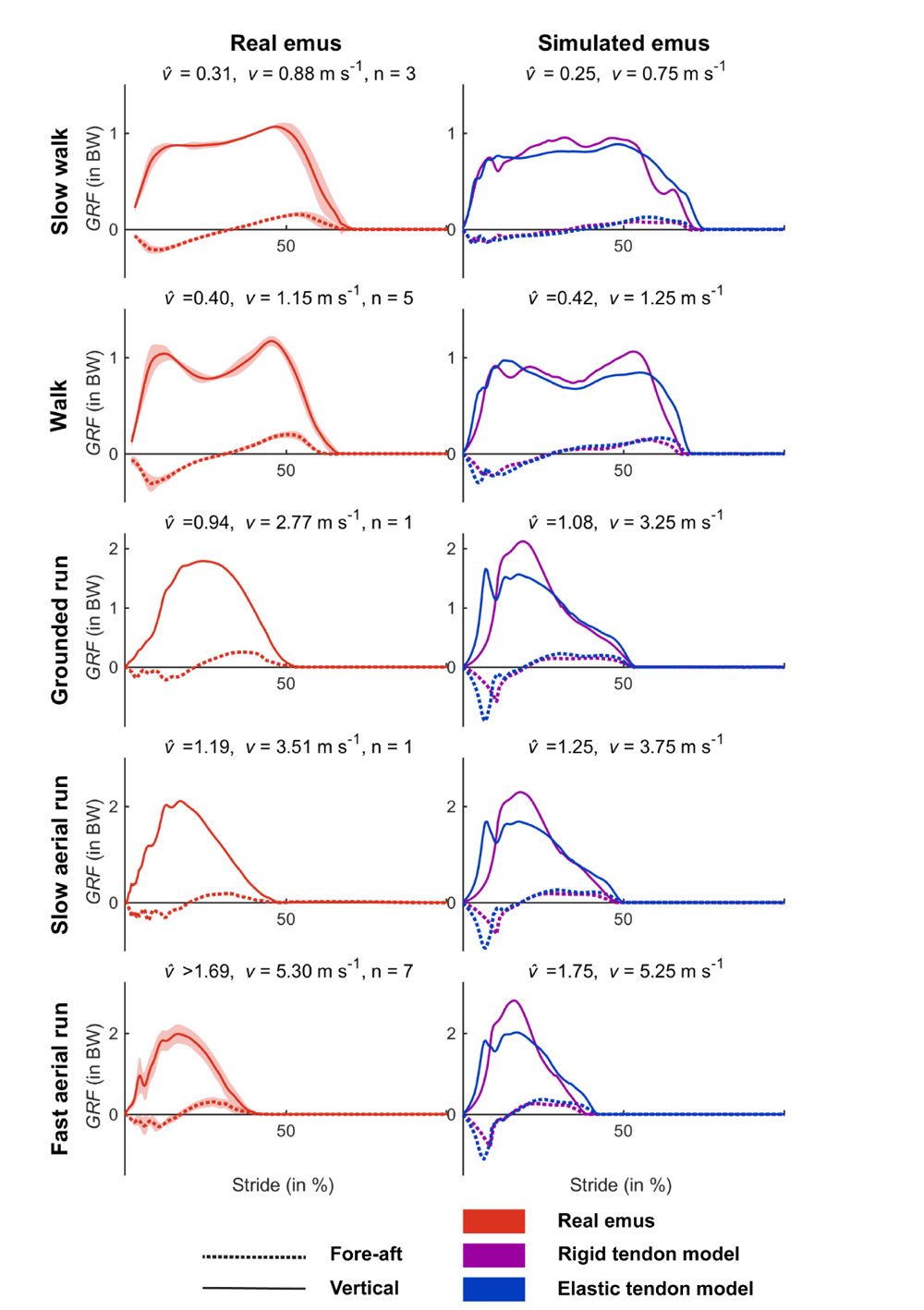
Ground reaction forces (normalized to bodyweight) of real emus (left) compared to our simulations (right) at dynamically similar speeds. These are the walk to run optimizations of the columnar model, optimizing for fatigue and *MCOT* simultaneously. The columnar model variants switched to grounded running at 3.25 m s^-1^, so the grounded running panel is at a different speed from the main manuscript and Figure S2. Two slow walk and all walk trials courtesy of J. Goetz. Grounded and slow aerial run trial, and one slow walk trial courtesy of J. Hutchinson. Fast aerial run trials courtesy of R. Main.

**Figure S2.**
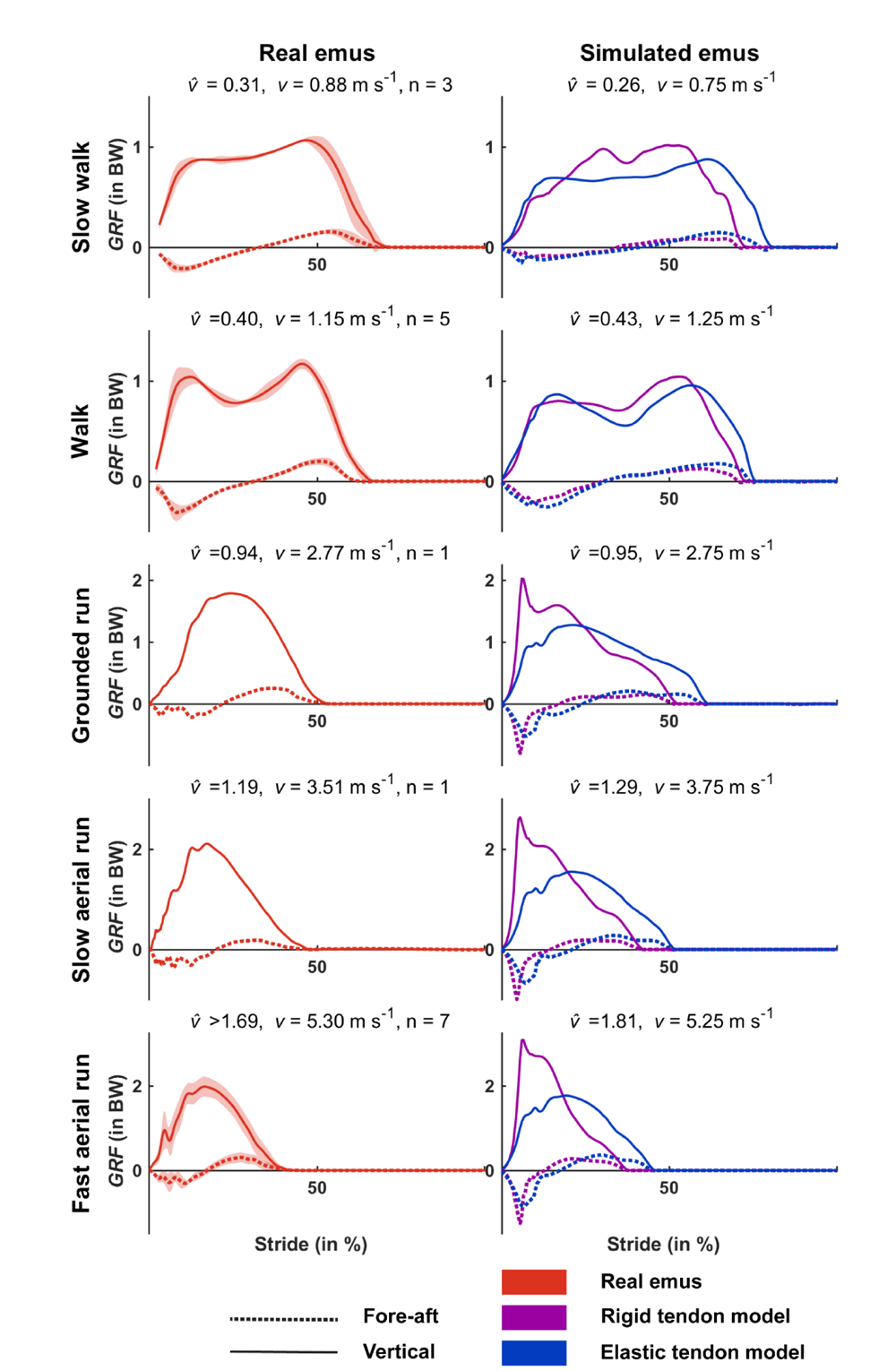
Ground reaction forces (normalized to bodyweight) of real emus (left) compared to our simulations (right) at dynamically similar speeds. These are the walk to run optimizations of the intermediate model, optimizing for fatigue and *MCOT* simultaneously. The rigid tendon model variant shows distinct impact peaks not present in real emus. Two slow walk and all walk trials courtesy of J. Goetz. Grounded and slow aerial run trial, and one slow walk trial courtesy of J. Hutchinson. Fast aerial run trials courtesy of R. Main.

**Figure S3.**
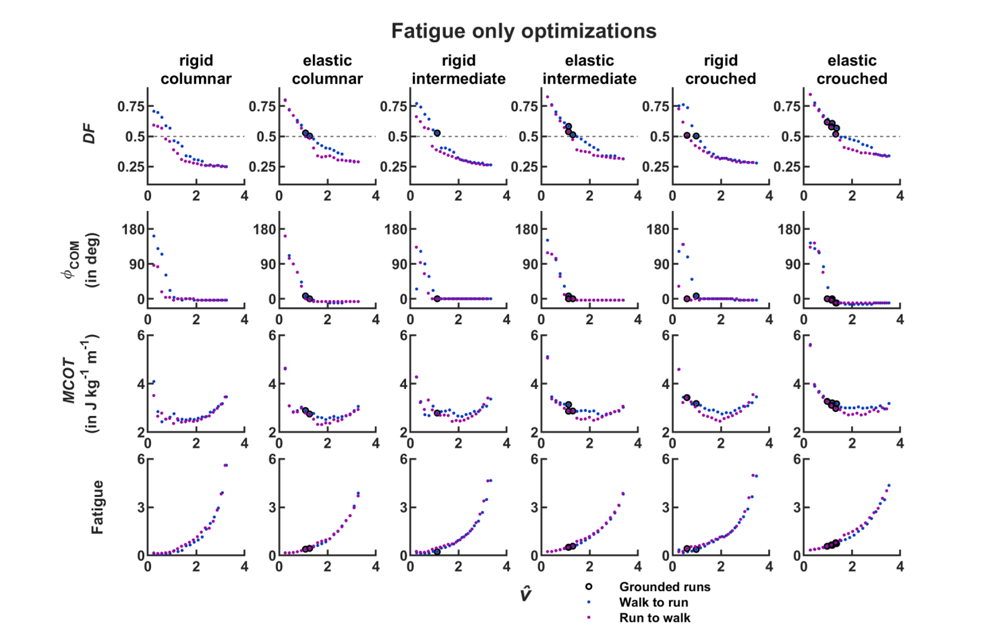
– Fatigue only optimizations. Each column represents a model iteration, with from top to bottom: Duty factors, phase angle of the *COM*, metabolic cost of transport, and fatigue. In all cases, we plot a speed range of 0.75 m s^-1^ – 9.75 m s^-1^, but differences in hip height affect relative speeds. Grounded runs are indicated separately. Similar to Figure 6 in the main manuscript, it can be seen that grounded runs occur more often from left to right, suggesting an effect of increasingly crouched postures, strengthened by the presence of elastic tendons. Fatigue is normalized and scaled (see methods).

**Figure S4.**
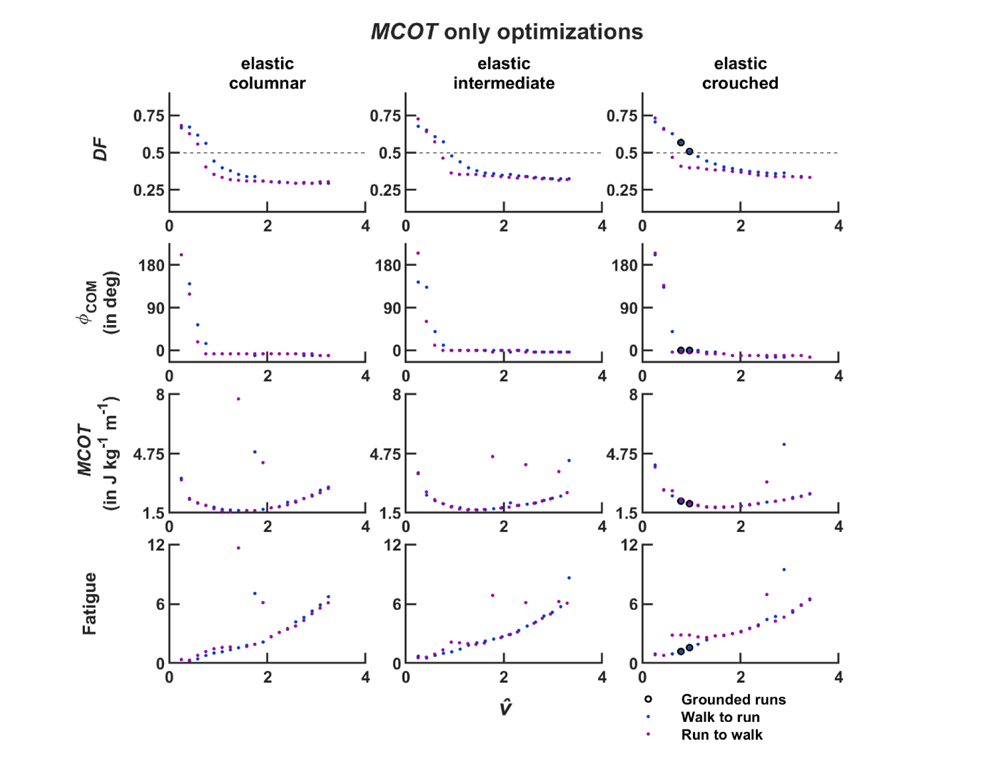
– *MCOT* only simulations. Each column represents a model iteration, with from top to bottom: Duty factors, phase angle of the COM, metabolic cost of transport, and muscle fatigue. From left to right: each model iteration. In all cases, we plot a speed range of 0.75 m s^-1^ – 9.75 m s^-1^, but differences in hip height affect relative speeds. Grounded runs are indicated separately. Similar to Figure 6 in the main manuscript, it can be seen that grounded runs occur more often from left to right, suggesting an effect of increasingly crouched postures. The rigid tendon models did not converge to single-period gaits when fatigue was excluded form the cost function, and are therefore not plotted. Fatigue is normalized and scaled (see methods).

**Figure S5.**
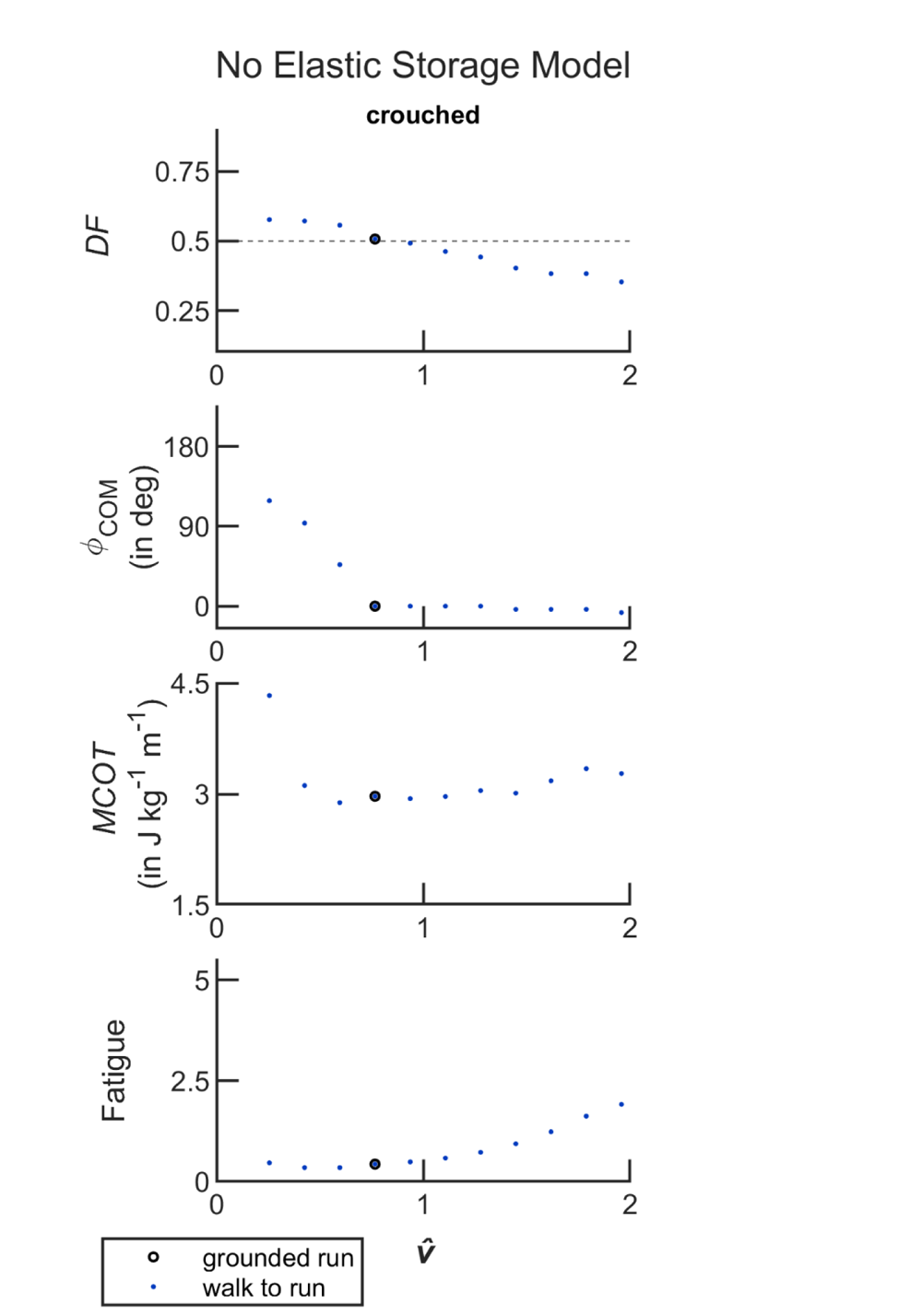
– Gait trials of the “no elastic storage” model variant. Top to bottom: Duty factors, phase angle of the COM, metabolic cost of transport, and muscle fatigue. Grounded runs are indicated separately. We plot a more limited speed range (0.75 – 5.75 m s^-1^), because this model variant only served to prove the possibility of grounded running in the absence of any elastic storage. Note that we rescaled the x-axis to accommodate this narrower range of speeds.

**Figure S6.**
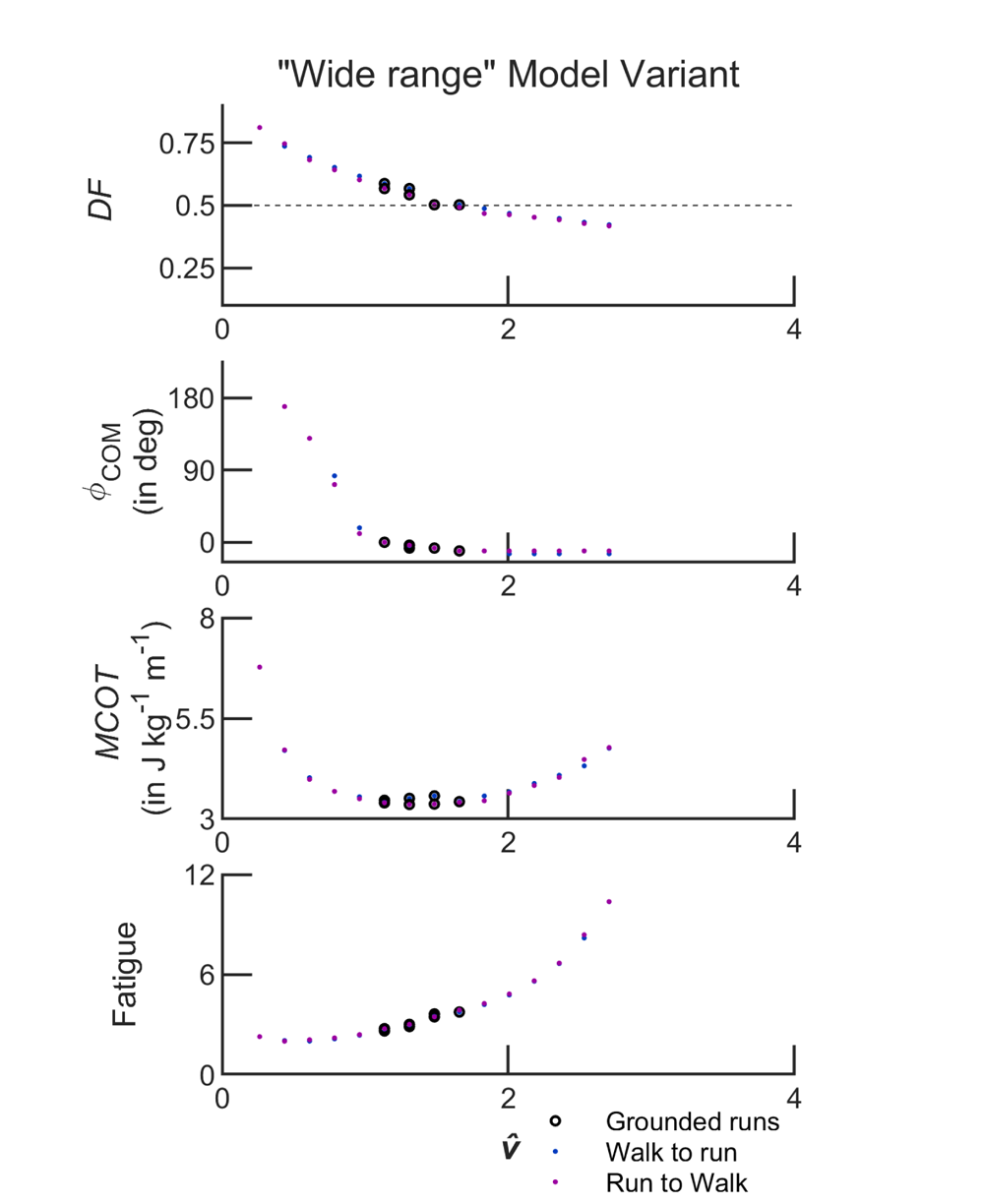
– Gait trials of the “wide range” model variant. Top to bottom: Duty factors, phase angle of the COM, metabolic cost of transport, and muscle fatigue. Grounded runs are indicated separately. We plot a speed range of 0.75 m s^-1^ – 7.75 m s^-1^. This model variant’s top speed was 8.25 m s^-1^. At 0.75 m s^-1^, only the “run to walk” trial converged, so we do not plot the other trial. Grounded runs are indicated separately. Similar to the crouched model variant, this model adopted grounded running over a wide range of speeds, in both optimization directions. Fatigue is normalized and scaled (see methods). Both *MCOT* and Fatigue were much higher in this model variant, and as such we rescaled their respective y-axes with respect to the figures in the main manuscript.

**Figure S7.**
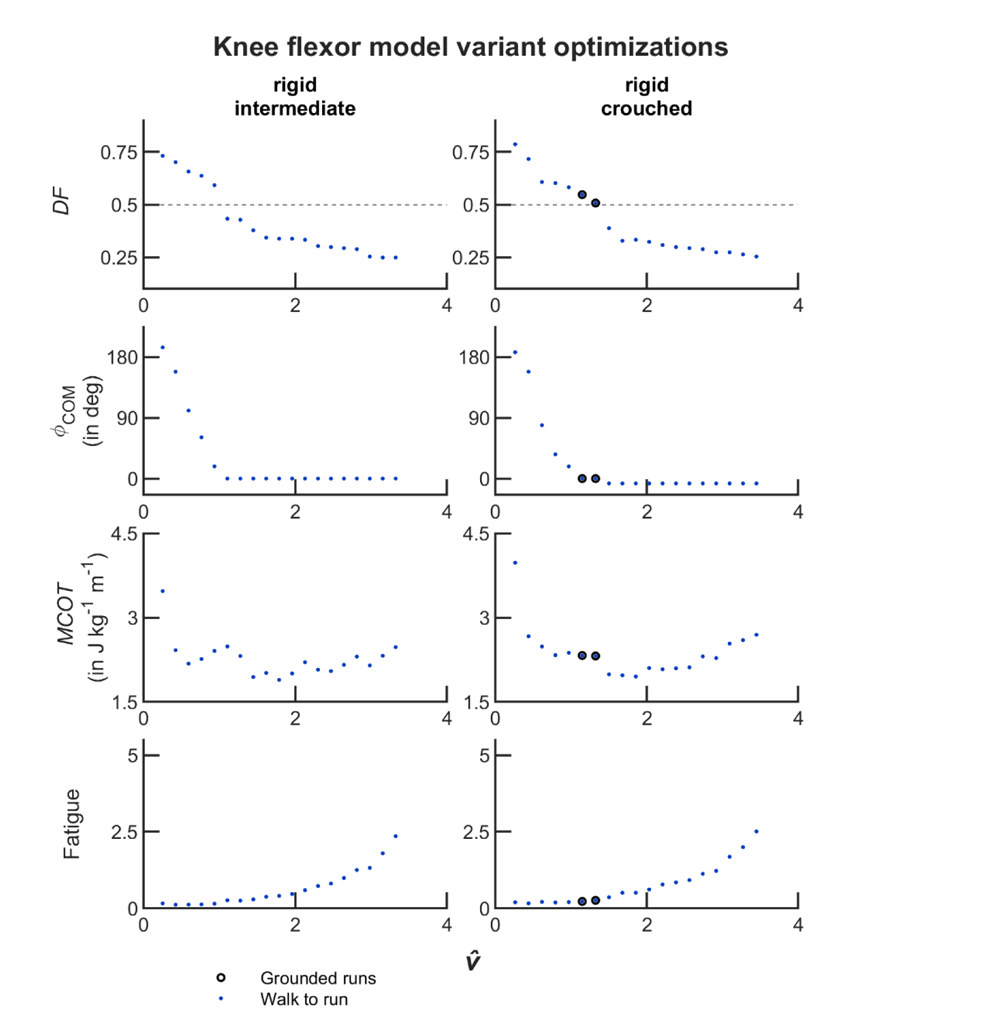
– Gait trials of the model variant with a mono-articular knee flexor, using intermediate and crouched tuning (left and right columns, respectively). Top to bottom: Duty factors, phase angle of the COM, metabolic cost of transport, and muscle fatigue. Grounded runs are indicated separately. It can be seen that the crouched model adopted grounded runs, whereas the intermediate did not. We plot a speed range of 0.75 m s^-1^ – 9.75 m s^-1^.

**Figure S8.**
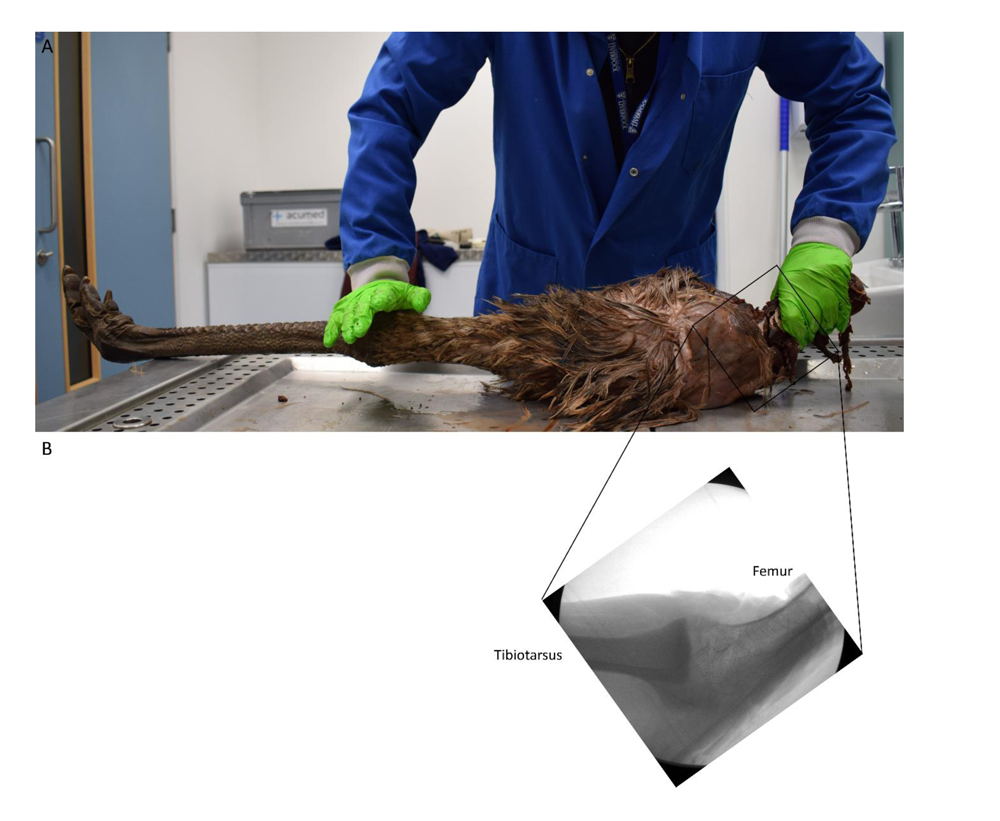
– **Maximal knee extension in the knee of the emu.** (**A**) Photograph in lateral view of a right leg. The experimenter is exerting a substantial amount of force on the femur and the tibiotarsus to ensure the knee is in maximal extension, this is likely more external extension moment than the knee experiences in vivo. Although the exact angle can only be estimated, it is clear that the knee cannot fully extend. (**B**) Radiograph of the same knee, lateral view. The radiograph has been mirrored to match the view in panel (A). The knee was wedged into full extension in a box, but the external extension moment was likely less than in panel (A). In this setup, the angle between the femur (left) and tibia (right) was approximately 138°, suggesting that maximal knee extension in the emu is ∼42° less than perfect alignment between the femur and the tibiotarsus. In our model, this corresponds to a knee angle of −25°.

## Supplementary Texts

### Muscle mass

Lamas et al. (*33*) presented two datasets of emu muscle masses, which we will refer to as the L-UK (n = 17) & and L-US (n = 29) datasets. Goetz et al. (*49*) also report muscle masses for a single specimen (“emu #3” from their study). The original study did not report the body mass of this specimen, but dr. Goetz reported its body mass as “95 lbs (43.1 kg)” in a supplementary dataset shared with us. These datasets were collected by different investigators who used different muscle-nomenclature. The L-UK dataset uses nomenclature that is similar to that the Handbook of Avian anatomy (*77*), and myology work in ostriches (*86*, *87*). The L-US dataset & Goetz et al. (*49*) used a different nomenclature (citing Patak & Baldwin (*19*)), which in some cases changes the interpretation of the muscle names (although the situation in ostrich myology was slightly different, see the appendix in Hutchinson et al. (*51*) for a discussion that is relevant here).

Similar to (*33*, *51*), we adopted the nomenclature proposed by (*77*), and thus converted the nomenclature of the L-US & Goetz et al. 2008 datasets. We predominantly used the muscle masses as a guide to confirm that these reinterpretations were consistent. We cross-validated these decisions by examining the photos and descriptions in the PhD thesis of A. Koenders (Patak) (*21*), which later led to the anatomical redescription of Patak & Baldwin (*19*). The most important difference is that Patak & Baldwin recognize an iliofemoralis cranialis (IFCR) (*19*) – this muscle is referred to as iliotrochantericus caudalis in the Handbook of Avian Anatomy (*77*), and is adopted in (*33*, *51*, *87*). A further difference is that Patak & Baldwin adopt the name ilioischiofemoralis (*19*) for a muscle that is more commonly referred to as caudofemoralis pars pelvica (*33*, *51*, *77*).

### Specifically, we converted

iliofemoralis cranialis (IFCR) to iliotrochantericus caudalis (ITC);

iliotrochantericus caudalis (ITC, only in L-US) to iliotrochantericus medius (ITM).

ilioischiofemoralis (IIF) to caudofemoralis pars pelvica (CFP);

femorotibialis externus (FTE, both heads) to femorotibialis lateralis (FMTL);

femorotibialis medius (FTM, only in L-US) to femorotibialis intermedius (FMTIM);

femorotibialis internus (FTI, all three heads) to femorotibialis medialis (FMTM);

Goetz et al. (*49*) do not report the same number of digital flexors (they do not report flexor perforatus digiti III & IV), and are the only group to report a measurement for plantaris, which other authors report as ‘ligamentous’ (*51*). We could not find a clear correspondence between their digital flexor measurements and those of Lamas et al. (*33*). We suspected conflicting identifications of muscles, and thus excluded the Goetz. et al measurements of digital flexors (PLANT, PPII, PPIII, PII, their abbreviations). Similarly, the L-US dataset seemed to have some clear outliers (Lamas et al. attribute discrepancies in the dataset to a different study goal when the L-US dataset was initially gathered (*33*)). We excluded all US measurements of TC, two US measurements of ITM, and one US measurement of ILPO. Goetz et al. reported some muscles summed together (FCLP & FCLA, and OMII & OMIP). These masses were split up according to the ratios in the Lamas dataset. L-US only included measurements for FCLP, but based on the relative muscle masses in the L-UK dataset, we interpret these measurements as FCLP+FCLA, and thus also split them up.

To visualize the ontogenetic effects, we first sorted all the specimens in ascending order of body mass (range: 3.6 – 51.7 kg). Next, we plotted the average relative hindlimb muscle mass (total hindlimb muscle mass / body mass) against number of specimens excluded (Figure ST 1, left). The figure shows a portion of roughly linear increase (when excluding light/young specimens), after which the average muscle mass remains roughly stable (unless too many specimens are excluded). We fit two straight line segments through these data, and the sum of squared differences was minimal when the inflection point was set at 17 excluded specimens (Figure ST 1, right). This can be interpreted as a lower bound on number of specimens to exclude, which would result in the lightest included specimen being merely 11.1 kg. To be slightly more conservative, we excluded two more specimens (19 in total).

**Figure ST1.**
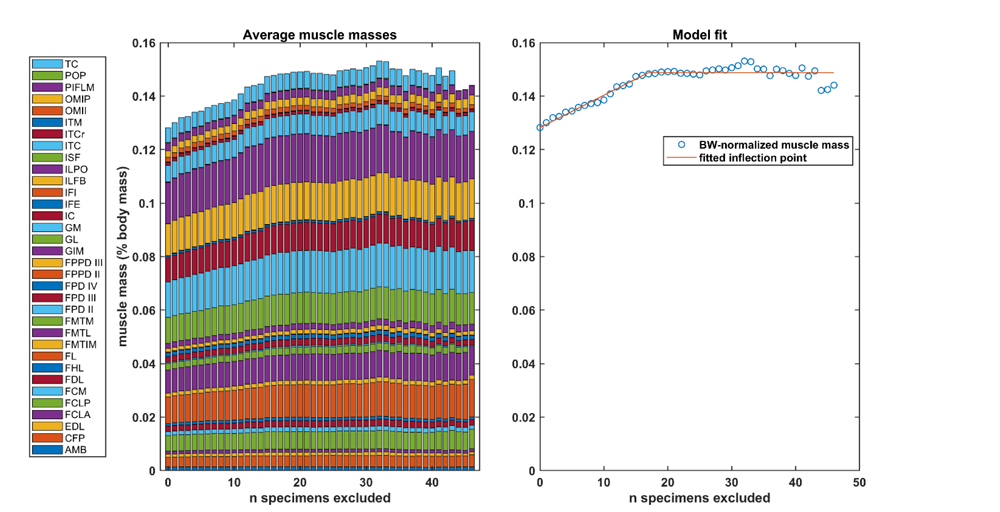
Left: Average muscle masses, normalized to body weight, sorted per muscle. See Table 2, main manuscript, for muscle abbreviations. On the x axis, we plot the effect of excluding specimens (sorted by ascending body mass) when computing the average. It is clear that ontogenetic differences in total muscle mass affects the average, when the lighter specimens are included. Right: We fit two simple line segments (linear increase, followed by stable muscle mass). The sum of least squares was minimal when the inflection point was at 17 specimens excluded (we plot this line). This represents a lower bound on the number of specimens to exclude due to ontogenetic effects. We excluded 19 specimens to be conservative.

### Muscle function mapping

Iliotibialis lateralis has two distinct sections with opposing functions about the hip: a pre-acetabular part (ILPR, hip flexor knee extensor), and a post-acetabular part (ILPO, hip extensor knee extensor). Goetz et al. were the only investigators who reported masses for these heads separately. In their data, ILPR comprised 29% of the total mass, and we used this ratio to map the mass of iliotibialis lateralis onto the two functions.

There seems to be a disagreement on the (sagittal plane) functions of Ambiens (AMB) across both the hip and the knee (which is also true for other ratites, e.g., (*51*)). It is predominantly a hip adductor, its flexion or extension function is secondary – and since it inserts on the medial side of the knee (*19*, *33*), its function across the knee is also uncertain. We have opted to distribute its mass across all three bi-articular hip functions.

Similarly, Femorotibialis medialis (femorotibialis internus), appears to have a knee-adduction function, and may secondarily act as a mono-articular knee-flexor. The model iterations used for our main analysis do not include this muscle, we added its mass to the hip-extensor knee-flexor (HEKF). We have included a sensitivity analysis to show that its inclusion results in similar gaits.

### Kinetic/kinematic data processing

Main manuscript Figure 4, the *GRF* comparison figure, is based on data from three different studies, kindly shared with us by Jessica Goetz (from (*49*)), John Hutchinson (from (*26*, *85*)) and Russel Main (from (*17*)).

Slow walk: average of 2 trials from Goetz. et al., and one from Lamas / Bishop et al.

Walk: average of 5 trials from Goetz et al.

Grounded run: Single trial from Lamas / Bishop et al.

Slow aerial run: Single trial from Lamas / Bishop et al.

Fast aerial run: average of 7 trials by Main et al.

The data reported in Goetz et al. 2008 (*49*) was cropped to the stance phase. J. Goetz kindly shared their original dataset of two emus (5 trial per individual) with us (*GRF* data low-pass filtered at 15 Hz and down sampled to 6 Hz, kinematic marker data filtered at 6 Hz, and corresponding videos of the trials). By identifying gait events initial contact and toe-off for both legs in the original video files, we estimated full stride periods and duty factors (assuming bilateral symmetry). We calculated corresponding forward velocities from the raw marker data. We calculated *h* for this dataset as the average hip height (from the kinematic data) over all trials for each individual emu. We excluded two trials for Emu #2 (E2C1T5) because it did not walk in a straight line. In total, this gave us eight extra datapoints for *L^* and *DF*. Based on the *GRF* traces, we excluded one more trial suspecting a movement asymmetry.v The numerical value of Goetz’s processed *GRF* data seem to be scaled by approximately a factor of two (see also Figure 2 in (*49*), which is not dynamically consistent with a walking gait). We enforced dynamic consistency by ensuring the average vertical *GRF* of one foot was equal to body weight, similar to (*26*).

John Hutchinson kindly provided three *GRF* timeseries and corresponding kinematic marker trajectories of emus. We estimated the stride time from contralateral foot contact in the *GRF*, and normalized the forces to body weight. Hip height was already known from (*26*). We recomputed *v* from the marker data. *GRF* sample frequency was 1000 Hz, and we plot the unfiltered data. Kinematic data was collected at 250 Hz.

Russel Main kindly provided videos, *GRF* data, and corresponding body masses of four emus (body mass 41.8 – 51.7 kg, three trials per emu). We estimated the stride times from the video files (collected at 250 Hz). Hip heights of the emus during walking speed were unavailable. By assuming a hip height of 1 m (an overestimate, based on comparisons with (*26*)), we acquired a lower bound on *v^*.

